# Complexities of recapitulating polygenic effects in natural populations: replication of genetic effects on wing shape in artificially selected and wild caught populations of *Drosophila melanogaster*

**DOI:** 10.1101/2022.05.12.491649

**Authors:** Katie Pelletier, William R. Pitchers, Anna Mammel, Emmalee Northrop-Albrecht, Eladio J. Márquez, Rosa A. Moscarella, David Houle, Ian Dworkin

## Abstract

Identifying the genetic architecture of complex traits is important to many geneticists, including those interested in human disease, plant and animal breeding, and evolutionary genetics. Advances in sequencing technology and statistical methods for genome-wide association studies (GWAS) have allowed for the identification of more variants with smaller effect sizes, however, many of these identified polymorphisms fail to be replicated in subsequent studies. In addition to sampling variation, this failure to replicate reflects the complexities introduced by factors including environmental variation, genetic background, and differences in allele frequencies among populations. Using *Drosophila melanogaster* wing shape, we ask if we can replicate allelic effects of polymorphisms first identified in a GWAS (Pitchers et al. 2019) in three genes: *dachsous (ds)*, *extra-macrochaete (emc)* and *neuralized (neur),* using artificial selection in the lab, and bulk segregant mapping in natural populations. We demonstrate that multivariate wing shape changes associated with these genes are aligned with major axes of phenotypic and genetic variation in natural populations. Following seven generations of artificial selection along the *ds* shape change vector, we observe genetic differentiation of variants in *ds* and genomic regions containing other genes in the hippo signaling pathway. This suggests a shared direction of effects within a developmental network. We also performed artificial selection with the *emc* shape change vector, which is not a part of the hippo signaling network, but showed a largely shared direction of effects. The response to selection along the *emc* vector was similar to that of *ds*, suggesting that the available genetic diversity of a population, summarized by the genetic (co)variance matrix (**G**), influenced alleles captured by selection. Despite the success with artificial selection, bulk segregant analysis using natural populations did not detect these same variants, likely due to the contribution of environmental variation and low minor allele frequencies, coupled with small effect sizes of the contributing variants.

## Introduction

Dissecting the genetic architecture underlying complex traits remains challenging, because of the joint contributions of many alleles of small effect, genotype-by-environment interactions, and other factors. Progress in sequencing technology in conjunction with development of GWAS statistical methodologies has enabled identification of loci contributing to numerous complex traits and diseases. However, such mapping approaches identify only a subset of loci contributing to trait variation (Visscher et al., 2017). In part, this reflects the low power to detect rare alleles, and those with small effects (Tam et al., 2019). For alleles that are relatively common in a population, replication rates between GWAS studies are high, even when effect sizes are small (Marigorta et al., 2018). However, GWAS studies have failed to replicate the effects observed in many candidate gene studies, in part due to the fact that many alleles identified in these studies are rare in populations, and require very large cohorts to detect (Fritsche et al., 2016; Ioannidis et al., 2011).

In cases where an association is replicated between studies, the magnitude of the effect can vary substantially between different cohorts or populations (CONVERGE consortium, 2015; Marigorta et al., 2018). Differences can arise because of genetic background due to epistatic gene by gene (GxG) interactions, or due to gene-by-environment (GxE) interactions. The initial estimates of effect size will be biased upwards if statistical testing in the initial cohort is used to determine which SNPs are chosen for replication studies. It is important to understand which of these causes of differences in effect size are of practical significance when we want to generalize results to different populations or environments.

In this study, we focus on the issue of replication in a multivariate context, where the joint inheritance of multiple features are simultaneously investigated. We will refer to the suite of measured features as a ‘multivariate trait’ for convenience. In this case, what we want to estimate is the vector of effects of each SNP on all measured features. Each SNP may have a unique combination of effects. Univariate effects vary only in magnitude, as we can only infer effects on a single feature. For a multivariate trait, estimated genetic effects vary in magnitude, the sum of effects on all traits, and also in direction, how the total effect is allocated among different features (Melo et al., 2019). The ability to study the direction along with the magnitude of genetic effects provides an additional and important way of assessing repeatability. For a univariate trait, there is a 50% chance that the replicate estimate will be in the same direction as the original estimate, even with no true effect. By contrast, the probability of a “replicated” genetic effect sharing a similar direction by chance alone decreases as the number of measured features increases (Marquez and Houle, 2015; Stephens, 2013).

Studying genetic effects in a multivariate context is beneficial in other ways. First, it has been demonstrated both empirically and via simulations, that genetic mapping for multivariate traits generally increases statistical power over trait by trait analyses (Fatumo et al., 2019; Pitchers et al., 2019; Porter and O’Reilly, 2017; Shriner, 2012). Second, some multivariate traits cannot be sensibly reduced to a single measurement. The wing shape we study is a great example of such a multivariate trait. We have good reason to believe that wing shape is important for flight (Ray et al., 2016), but we cannot yet say that any feature, such as wing length or width, is more or less important than any other. Natural selection on wing shape may affect any or all combinations of measurements.

Perhaps most importantly, traits are not inherited in isolation, but are the joint outcome of an integrated developmental process that results in extensive genetic correlations that can have important effects on evolution. The main source of such correlations are the patterns of pleiotropic effects generated by mutational effects. Multivariate studies of inheritance allow pleiotropic effects to be estimated in a rigorous and justifiable manner (Melo et al., 2019). The multivariate breeder’s equation, ***Δ***𝑧 = 𝐺𝜷, enables short term prediction of evolutionary responses. Key to understanding how populations respond to selection in the short term requires an understanding of properties of the genetic (co)variance matrix (**G**), and in particular the axis of greatest genetic variation, **g**_max_. Studies demonstrate that the direction of **g**_max_ influences evolutionary trajectories (Blows and McGuigan, 2015; McGuigan, 2006; Schluter, 1996). The degree to which genetic effects associated with particular variants align to major axes of genetic (co)variance, expressed through **G**, may provide insights into which alleles are most likely to be “captured” by selection (Pitchers et al., 2019).

Due to the polygenic nature of complex traits, including multivariate ones, it is important to consider not only the direction of effect for alleles in a single gene but also correlated effects between genes contributing to the phenotype. Interestingly, initial comparisons of directions of genetic effects among induced mutations in two *Drosophila melanogaster* wing development pathways showed only partially correlated effects on wing shape within and between pathways (Dworkin and Gibson, 2006). However, recent work has demonstrated that despite large differences in magnitude, the direction of genetic effects of variants segregating in populations are sometimes similar to those from validation experiments using RNAi knockdown of those same genes (Pitchers et al., 2019). Additionally, Pitchers et al. (2019) demonstrated this shared direction of effect could also be shared between a SNP and RNAi knockdown of other genes in the same signaling pathway, such as those involved with hippo signaling, a key pathway involved with wing growth and morphogenesis (Pan et al., 2018).

Pitchers et al. (2019) identified over 500 polymorphisms contributing to wing shape variation in the *Drosophila* genetic resource panel (DGRP). Among these, the hippo pathway was over-represented in SNPs associated with wing shape (Pitchers et al., 2019). The degree to which identified hippo pathway variants reflect allele specific effects, differences in magnitude of genetic effects, and even the large statistical uncertainty associated with genetic effects of small magnitude are unclear. Given common dominance patterns, and the likely non-linear genotype-phenotype relationships of most genetic effects, small to moderate changes in gene function may result in modest phenotypic effects (Green et al., 2017; Melo et al., 2019; Wright, 1934). Large effect mutants and many RNAi knockdown studies have moderate to large phenotypic effects that are not reflective of the magnitude of genetic effects of SNPs contributing to phenotypic variance in natural populations.

The expression of genetic effects also depends on genetic and environmental context, with gene-by-gene (GxG) and gene-by-environment (GxE) interactions contributing to phenotypic variation. The context-dependence of genetic effects for a multivariate trait has been demonstrated for *Drosophila* wing shape. Variants in *Epidermal growth factor receptor* (*Egfr*), influencing *Drosophila* wing shape are replicable in both lab reared, and wild-caught cohorts (Dworkin et al., 2005; Palsson et al., 2005; Palsson and Gibson, 2004). However, in replication studies, effect sizes of alleles were diminished in both outbred populations and wild cohorts. In the latter case the same variant explained 1/10 of the phenotypic variance explained in the initial study (Dworkin et al., 2005). Interestingly, in a series of experimental crosses among strains, the effects of the SNP were replicable for direction and magnitude in multiple experimental assays and crossing schemes. Despite this, the genetic effect on wing shape from this SNP largely disappeared in one natural population (Palsson et al., 2005). A number of reasons have been proposed for the failure to replicate genetic effects including environmental effects, differences between controlled lab and natural environments (Dworkin et al., 2005), and genetic background (Greene et al., 2009), among others. Because both environment and genetic background likely affect the genotype-phenotype map in a non-linear fashion (Wright, 1934), it is important to test observed associations in other experimental contexts.

A promising approach to confirm the estimated effects of candidate genetic variants is to test whether they respond to artificial selection in the direction of the inferred effect. This approach is particularly relevant to evolutionary questions, but has rarely been used. In this study, we use artificial selection and bulk segregant analysis (BSA), to replicate and validate SNPs associated with three genes, previously identified in a GWAS of *Drosophila* wing shape (Pitchers et al., 2019); *dachsous (ds), an atypical cadherin involved with hippo signaling;* the transcriptional co-repressor *extra-macrochetae (emc),* and the E3 ubiquitin ligase *neuralized (neur)*, involved with Notch signaling. Using the vectors of shape change based on RNAi knockdowns of each gene, we demonstrate that the direction of shape change for these genetic effects is aligned with major axes of natural phenotypic and genetic variation. Using artificial selection based on the direction of shape change defined by RNAi knockdown, we were able to replicate the effects observed for *ds,* but not *emc*, likely due to the available genetic diversity in the population. We then asked if these effects could be replicated in a natural population using a bulk segregant approach, observing little evidence for replication in these samples. We discuss our results in the context of the replicability of genetic effects and the shared direction of genetic effects due to shared developmental processes.

## Methods

### Source Populations and phenotypic analysis

#### Drosophila strains

Phenotype data for the *Drosophila* genetic resource panel (DGRP) was collected for 184 strains as part of a GWAS study as described in Pitchers *et al* (2019). Genotype data for these strains was obtained from freeze 2 of the DGRP (Huang et al., 2014). For replication using artificial selection, 30 DGRP strains were used: DGRP-149, 324, 383, 486, 563, 714, 761, 787, 796, 801, 819, 821, 822, 832, 843, 849, 850, 853, 859, 861, 879, 887, 897, 900, 907, 911, 913. These strains were selected to increase genetic variation at the *ds* locus (Supplemental Figure 1, Table 1). Reciprocal pairwise crosses between the 30 selected DGRP strains were used to create heterozygotes and these 30 heterozygous genotypes were successively pooled for 4 subsequent generations, allowing for recombination. After pooling, the synthetic outbred population was maintained for approximately 47 subsequent generations (allowing for recombination) before the start of artificial selection experiments.

**Table 1.**
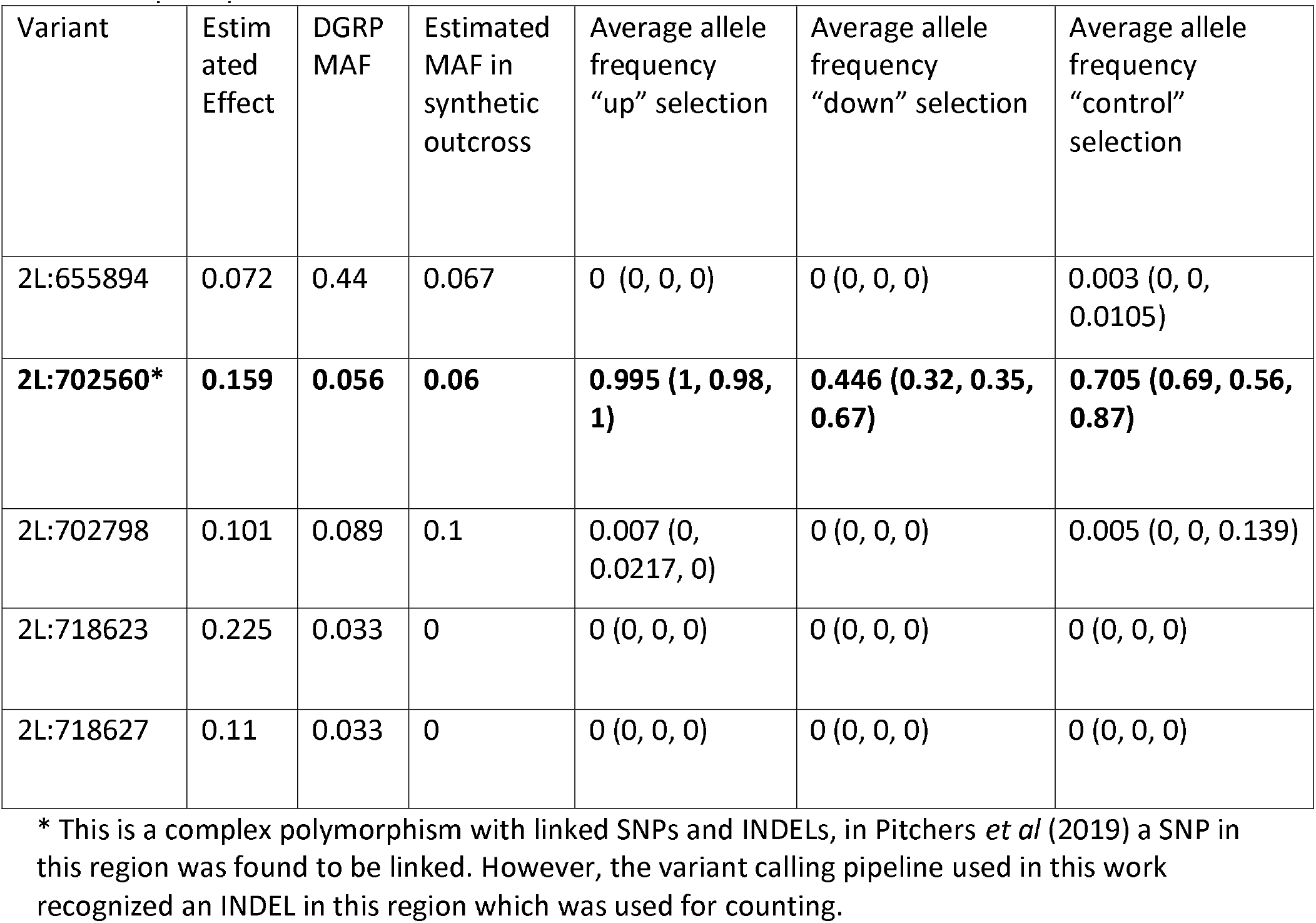
Variants from Pitchers et al. (2019) in ds artificial selection experiment. Estimated effect sizes for SNPs are estimated from a GWAS in the DGRP using LASSO regularized coefficients. Average frequency is given with replicate lineage frequencies in brackets. Estimated effect is the *l^2^-norm* of shape differences associated with the variant. MAF = minor allele frequency.

For the replication in wild-caught populations using BSA, individuals were collected via sweep-netting from orchards and vineyards in Michigan and after species identification, stored in 70% ethanol. In 2013 and 2014, cohorts were collected from Fenn Valley Winery (FVW13 and FVW14 respectively, GPS coordinates: 42.578919, −86.144936). Additionally in 2014, cohorts were collected from Country Mill Orchard (CMO, GPS coordinates: 42.635270, −84.796706), and Phillip’s Hill Orchard (PHO, GPS coordinates: 43.117981, −84.624235). For all collected cohorts, except for the FVW14 collection, only males were used in this study given difficulties distinguishing *Drosophila melanogaster* and *D. simulans* females morphologically. For the genomic analysis of the FVW14 wild caught population (below) we utilized both males and females as the number of individuals was insufficient otherwise. For the collection where females were included in the study, there is no evidence of contamination with*D. simulans* as all dissected wings were classified as *D. melanogaster* using linear discriminant analysis (LDA). LDA was trained using male wings from the collected *D. melanogaster* data set and males from *D. simulans*. There was 100% agreement between the classification of females within each species with our phenotypic classification, indicating that it is unlikely that *D. simulans* females were included in our samples (Supplemental Figure 2).

#### Morphometric Data

Landmark and semi-landmark data were captured from black and white TIFF images using the pipeline described in Houle et al. (2003). First, two landmark locations, the humeral break and alula notch, were digitized using tpsDig2 (version 2.16). Wings (Van der Linde 2004– 2014 , v3.72) software was used to fit nine cubic B-splines, and manually correct errors. All shape data was subjected to Procrustes superimposition (registration), removing the effects of location, isometric scaling, and and minimizing effects of rotation, , via an iterative least squares approach (Rohlf and Slice, 1990). Generalized Procrustes superimposition (registration) and extraction of 14 landmarks and 34 semi landmarks was done using CPR v1.11 (Márquez 2012–2014, Figure 1). Sliding of semi-landmarks utilized minimization of Procrustes Distance as the objective function. Superimposition results in the loss of 4 possible dimensions of variation while semi-landmarks are constrained to vary along one “axis”, restraining these points to approximately a single dimension of variation each. This results in a total of ∼58 available dimensions of shape variation, that can be summarized using the first 58 Principal components (PCs). Allometry was adjusted for in the analysis by fitting a model for landmark coordinates onto centroid size, and using the residuals from this model (Klingenberg, 2022). By accounting for the allometric component of shape, shape variation associated with size variation can be accounted for (Supplemental Figure 3). For most analyses, ‘allometry corrected’ shape data were used, with the exception of shape models fit using the Geomorph package in R, where Procrustes landmarks were used and centroid size was included as a predictor in the model.

**Figure 1.**
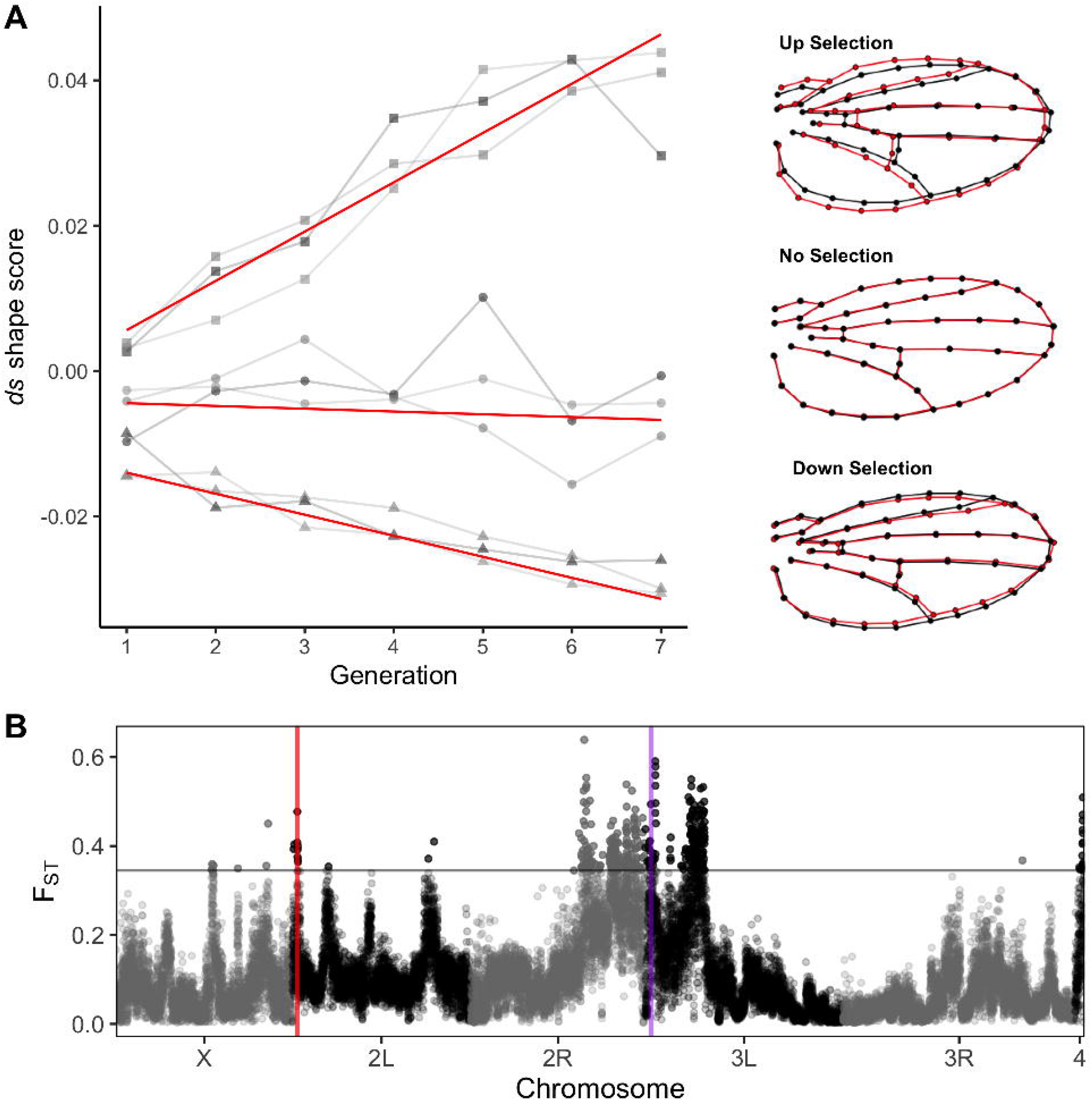
Projections of data onto RNAi shape change vectors are correlated with major axes of shape variation among DGRP strains. (A) Shape change vectors from RNAi titration experiments for *ds, emc* and *neur* were used, and DGRP line means were projected onto these vectors to calculate shape scores. Eigenvectors for the PCA were estimated based on the same DGRP line means. Vector correlations between shape change vectors from RNAi knockdown:*ds* − *emc*: 0.65, *ds* − *neur*: 0.03, *emc* − *neur*: 0.30. (B) Effect of *ds* shape change estimated from RNAi knockdown, effects not magnified. (C) Landmarks (red) and semi-landmarks (blue) used in geomorphic morphometric analysis on a *Drosophila* wing. PCs 1−3 account for 22%, 20% and 9% of the overall, among DGRP shape variance.

#### Generation of shape vectors for artificial selection and bulk segregant analysis

A panel of shape change vectors was estimated using the progesterone-inducible Geneswitch GAL4, under the regulation of an ubiquitous *tubulin* driver, to drive the expression of RNAi for genes of interest (*ds*, *emc*, *neur*), as previously described in Pitchers *et al.,* 2019. GAL4 expression was induced throughout larval development by adding mifepristone, an analog of progesterone, to the larval food. Knockdown was varied by assaying phenotypes at mifepristone concentrations of 0.3, 0.9, and 2.7 μM, plus a control without mifepristone. Wing shape change associated with knockdown of the gene of interest was estimated using multivariate regression of shape on concentration of mifepristone. Shape change vectors estimated from the RNAi experiments for *ds,emc* and *neur*, were used in this experiment (Figure 1b, Supplemental Figure 4). The magnitude (‘length’) of the vector measures how much shape change occurs per unit change in mifepristone. In general, vectors of greater magnitudes enable better estimate of direction of effect for shape change. As reported in Pitchers et al., (2019), the magnitude (*l*^2^*-norm*) of vectors for RNAi knockdown of these genes are 5.5 for *ds*, 2.8 for *neur*, and 0.44 for *emc*.

Shape data collected as part of a previous study (Pitchers et al., 2019) was used to assess the relationships between shape change vectors from the RNAi titration and **g**_max_, the first eigenvector of the **G** matrix estimated from DGRP line means. The effects of sex, centroid size and their interaction were removed using a linear model and these residuals were used to calculate shape score by projecting the data (see Supplemental Figure 5) onto the shape change vector estimated in each knockdown experiment. To assess major axes of genetic variation among DGRP strains, principal component analysis was performed on allometry adjusted model residuals (Supplemental Figure 5B). PCA was done in a similar manner for individuals from the wild caught cohorts. Correlations between the first three eigenvectors (“genetic PCs” including **g**_max_), the first three PCs from the wild caught cohorts and the shape scores for *ds*, *emc* and *neur* were calculated (Figure 1a, Supplemental Figure 5). From this, *ds, emc* and *neur* shape change vectors were selected for further experiments given high correlation with directions of natural genetic variation (Figure 1, Supplemental Figure 5). Note, as described below, while*ds* and *emc* were used for artificial selection, due to the similar response between them, we used *ds* and substituted *neur (*for *emc)* for the BSA.

#### Artificial selection of synthetic outbred population

The synthetic outbred population resulting from pooling DGRP lines was used as the parent population for artificial selection. Both the *ds* and *emc* artificial selection experiment were carried out with three independent replicates of each “up” and “down” selection regimes, along with unselected control lineages. Each generation, wings of live flies were imaged using the ‘wingmachine’ system and shape data collected (Houle et al., 2003, Van der Linde 2004– 2014 ,v3.72). Shape scores were calculated by projecting the data onto the *ds* or *emc* shape change vector as described above, and the 40 individuals each with highest or lowest shape scores, were selected to found the next generation (Supplemental Figure 5A). For the control lineages, 40 individuals were randomly selected for the next generation within each replicate lineage. Following seven generations of selection, 75 individuals from each lineage were selected for pooled sequencing, described below. The response to selection was evaluated both by computing Procrustes distance (PD) between average shape of wings between generations one and seven, and using shape scores (projections) with a linear mixed effect model allowing for the fixed effect factors of treatment and sex, continuous predictors of centroid size and generation, with third order interactions among these effects. The effect of generation was allowed to vary by replicate lineages (lmer(ds ∼ (CS + Sex + line + gen0)^3 + (1 + gen0|line:rep) ). Realized heritabilities were estimated separately for up and down selection lineages, from the slope of the regression of cumulative selection differentials on cumulative selection response, averaging over sex and with a random effect of replicate lineage.

#### Wild populations

For the BSA, wings for wild caught individuals were dissected and mounted in 70% glycerol in PBS. Images of wings were captured using an Olympus DP30B camera mounted on an Olympus BX51 microscope (Olympus software V.3,1,1208) at 20X magnification. When possible, both left and right wings were dissected, imaged and averaged to calculate an individual’s mean shape. For some individuals a wing was damaged so only one wing could be used. Shape was captured as described above. The total number of individuals phenotyped from each cohort can be found in Supplemental table 1.

To remove allometric effects in the data, shape was regressed onto centroid size and the model residuals were used for all subsequent morphometric analysis. Only data from males was used to compare shape in wild populations, although, including females from the FVW14 population and regressing shape onto centroid size and sex gave equivalent results (Supplemental Figure 6). To test for shape differences between collection cohorts, the effect of centroid size and collection cohort on shape were modeled (procD.lm(shape ∼ CS + pop_year)) using the procD.lm function in Geomorph v 3.1.3. (Adams and Otárola-Castillo, 2013) and distances between populations were calculated using the pairwise function. To select individuals for sequencing, a ‘shape score’ was calculated using the method described above.

Shape data was projected onto the vector of shape change defined by the *ds* or *neur* knockdowns. The *emc* projection vector was not used for BSA due to the high similarity with *ds* shape change (Figure 1), and the similarity of the selection response. Its inclusion would result in selection of largely the same cohorts of individuals for sequencing for both *ds* and *emc*. As an alternative, we utilized the *neur* shape vector as it was largely uncorrelated with that of *emc* and *ds*, but strongly correlated with natural variation in shape. The 75 most extreme individuals on the shape score distribution, within each wild-caught cohort, were selected for pooled sequencing. Allele frequencies within each population was estimated by sequencing 75 random individuals within each cohort. The difference vector between mean shapes of selected pools (within each population) was used to calculate Procrustes distance (PD) between pools and the correlation of this shape change vector with the selection vector used. An estimate of genetic distances between populations was calculated using allele frequencies (mapping pipeline described below) in the pools of the 75 randomly selected individuals using Bray’s distance with the vegdist() function from the vegan package (v2.6-2) in R.

#### Sequencing and Genomic Analysis

DNA extractions from pools of selected individuals was performed using a Qiagen DNeasy DNA extraction kit. Library preparation and Illumina sequencing was performed at the research technology support facility at Michigan State University. All library samples were prepared using the Rubicon ThruPLEX DNA Library Preparation kit, without a procedure for automatic size selection of samples. Paired end libraries (150bp) were sequenced using Illumina HiSeq 2500, with each sample (either one pool of 75 individuals in the BSA or one pooled replicate lineage in the artificial selection) being run on two lanes.

Reads were trimmed with Trimmomatic (v0.36) to remove adapter contamination and checked for quality using FastQC prior to alignment (Bolger et al., 2014). Trimmed reads were aligned to the *Drosophila melanogaster* genome (v6.23) using BWA-MEM (v0.7.8)(Li and Durbin, 2010). Sequencing replicates of the same biological samples were merged using SAMtools (v1.11). PCR duplicates were removed using Picard with the MarkDuplicates tool (v 2.10.3) and reads with a mapping quality score less than 20 were removed using SAMtools (Li et al., 2009). A local realignment around indels was performed using GATK using the IndelRealigner tool (v3.4.46). For artificial selection experiments, reads were merged for all up, down and control selection lines as replicates lineages were independent. For wild cohorts, pools were not merged between populations. mpileup files were created using SAMtools and used for subsequent genomic analysis. Highly repetitive regions of the*Drosophila* genome were identified and subsequently masked in mpileup files using RepeatMasker (v4.1.1) with default settings. INDELs and regions within 5bp of an indel were identified and masked using popoolation2 scripts. Population genetic statistics were calculated using PoPoolation (v1.2.2) and PoPoolation2 (v1.201) (Kofler et al., 2011b, 2011a).

For the BSA in the wild-caught cohorts, a modified Cochran-Mantel-Haenszel (CMH) test was used to measure significantly differentiated sites between pools of individuals. Sampling effects were accounted for using the ACER package (v.1.0) in R, assuming *N*_e_ *= 10*^6^ with 0 generations of differentiation between selected pools (Spitzer et al., 2020). To adjust for multiple testing, the p-value was corrected using a Benjamini-Hochberg correction (Benjamini and Hochberg, 1995) with an adjusted alpha of 0.05. For each significant site from the CMH test, using an adjusted p-value cut-off of 0.05, we identified the nearest gene using BEDtools (v2.19.1) (Quinlan and Hall, 2010). In addition, to account for sampling variation, we sampled genomic coverage to 75x for all samples, dropping sites that did not meet this threshold and repeating the CMH test. We confirmed that there was no association between genetic and shape differentiation between populations, and that the populations do not show strong phenotypic differentiation based on either overall shape variation, or shape scores used to identify selected individuals for BSA (Supplemental Figures 3 and 7). There was some variation among populations in overall wing size (Supplemental Figure 8), however we (assuming common allometry) adjusted for allometric effects on shape.

For artificial selection experiments, F_ST_ was calculated in 5000bp windows. We chose this window size as it is expected that blocks of LD in the synthetic outbred population will be much larger in comparison to that of the wild caught samples (King et al., 2012a; King et al., 2012b; Marriage et al., 2014). This statistic was used to compare the “up” selected pools to the “down” selected pools to help identify regions of differentiation between selected populations.

For the artificial selection comparisons, genes in regions of high F_ST_ were identified by finding overlaps between outlier windows and annotated *Drosophila* genes using GenomicRanges (v1.46.1) in Bioconductor. High F_ST_ was defined as F_ST_ values greater than three standard deviations above the mean. GO terms associated with identified genes were annotated using TopGO package (v2.34.0) (Alexa et al., 2006) in Bioconductor. GO enrichment was then performed to identify those terms overrepresented in the identified list using TopGO and a Fisher’s exact test. Over representation of 2 GO terms in outlier windows (hippo signaling, GO:0035329; negative regulation of hippo signaling GO:0035331) were tested using a permutation test that randomly sampled genomic windows from the total windows for which F_ST_ was calculated and the permutation was run 1000 times. The distribution of the ratio of observed to expected genes annotated with the term of interest within randomly sampled regions was compared to the number observed in the data.

#### Verification of *ds* indel in DGRP

Sanger sequencing was performed on individuals from a cross between DGRP lines predicted to have the polymorphism (DGRP 195, 28, 96, 48, 59, 801) and those without (DGRP 129, 301, 69, 385, 75, 83, 491, 34, 774) crossed to a line carrying a deletion in the region of interest (BDSC 24960) to account for potential residual heterozygosity in otherwise inbred strains. DNA was prepared by incubating flies in DNA extraction buffer (1mM EDTA, 25mM NaCl, 10 mM TrisHCl pH 7.5) for 10 minutes, followed by storage at −20 C. PCR application of the region of interest (Forward primer: ggagtacaagttgctcgaac: Reverse Primer: cagatcgtgttccctttagc) using Taq DNA polymerase (Gene DirectX) (PCR mix: 1uL DNA, 1uL forward primer, 1uL reverse primer, 0.5uL 10mM dNTPs, 2uL 10x PCR Buffer, 0.1 uL taq, H20 to 20uL). PCR conditions were as follows: 5 minutes 95°, (30 seconds 95°, 30 seconds 55°, 30 seconds 72°)x30. Reactions were checked on a gel and cleaned with the GenepHlow^TM^ Gel/PCR Kit (Geneaid). Sanger sequencing reactions were performed by the Mobix Lab at McMaster University. All alignments were created using ClustalOmega (Madeira et al., 2022).

#### Data Availability

All code and processed data needed to complete the analysis is available on GitHub at: https://github.com/DworkinLab/WingShapeBSA/. A static version of the repository is available on figshare (https://doi.org/10.6084/m9.figshare.22141154.v1).

All raw sequence data available as part of the NCBI Short Read Archive, BioProject PRJNA936488 (https://www.ncbi.nlm.nih.gov/bioproject/PRJNA936488/), with individual sequence accessions SAMN33354503 - SAMN33354634.

## Results

### *dachsous (ds)* shape change is aligned with major axes of genetic and phenotypic variation in natural populations

To assess the relationship between shape change vectors and axes of natural variation described in the the DGRP, mean shape vectors were calculated for each DGRP strain, then used in a PCA to summarize axes of variation among strains. Mean shape vectors for each strain of DGRP were projected onto shape change vectors for *ds emc*, and *neur*, defined from the RNAi knockdowns (see Supplementary Figure 5, which visually explains the procedure), generating gene specific “shape scores”. Correlations between shape scores for individual DGRP projected onto the shape change vectors (Figure 1, Supplemental Figure 5), and with PC1 generated from the DGRP (PC1_DGRP_) strains was estimated (PC1_DGRP_-*ds*: r = −0.56; PC1_DGRP_-*emc*: r = −0.45; Figure 1). The correlation of the DGRP data, projected onto each of the*ds* and *emc* shape change vectors was also correlated (Figure 1, *ds*-*emc*: r = 0.69). This is likely due to the correlation between gene specific shape change vectors themselves (r = 0.65), based on RNAi titration experiments. Projections of the DGRP data onto the vector defining the*neur* shape change is aligned with PC1 (PC_DGRP_-*neur*: r = −0.69) and PC3 (PC_DGRP_-*neur*: r = −0.64), indicating this as an important axis of shape variation in this population (Figure 1), that is moderately similar to projections onto *ds* (*ds*-*neur*: r = 0.56) and very similar to *emc* (*neur*-*emc*: r = 0.83) shape change vectors. Interestingly, the strength of the correlation for the DGRP strains projected onto these vectors, differs from the magnitude of correlations for the RNAi titration vector of *neur* with that of *ds* (r = 0.034) or *emc* (0.3). Because of these observed correlations, and previous associations observed (Pitchers et al., 2019) *ds, emc* and *neur* were selected as focal genes for subsequent studies.

We also examined the relationship between direction of phenotypic effects with the wild caught cohorts. For these samples, phenotypic variance for shape is due to the joint contribution of genetic and environmental effects. To illustrate the difference in shape variance in wild populations and the DGRP, we calculated correlations between the first three eigenvectors for shape in the DGRP, the combined wild cohorts as well as the CMO cohort alone. We observed low correlations between the DGRP eigenvectors and those estimated from wild populations (Supplemental Table 2). As observed with the DGRP, there is a substantial correlation between projections of shapes of individuals onto the *ds* shape change vector and PC1 (defined by phenotypic variation among wild caught files, PC_wild_) in most of the sampled cohorts (*ds*-PC1_wild_ PHO: r = 0.78; CMO: r = 0.87; FVW13: r = −0.22; FVW14: r = 0.95, Figure 2, Supplemental Figure 9). In cohorts where the *ds* shape change vector was not correlated with PC1, specifically the FVW13 collection, this vector is correlated with PC2 (*ds*- PC2_wild_ PHO: r = 0.12; CMO r = −0.44; FVW13: r = −0.63; FVW14: r = 0.19; Figure 2; Supplemental Figure 5). The pattern for the wing shape from wild-caught individuals projected onto the *emc* shape change vector was generally similar to that observed for *ds* (Figure 2). We also observe a correlation between *neur* shape change and PC1 in most cohorts (*neur*-PC1_wild_ PHO: r = 0.51; CMO: r = −0.051; FVW12: r = −0.95 FVW13; FVW14: r = −0.084; Figure 2; Supplemental Figure 7). As with the *ds* shape change vector, in some cohorts such as the CMO the stronger correlation is between the *neur* shape change vector and PC2 (PHO: r = 0.22; CMO: r = −0.57; FVW13: r = −0.0.059; FVW14: r = 0.85; Figure 2; Supplemental Figure 7). Interestingly, in the CMO cohort, the correlations between the projection of shape data onto the*ds* and *neur* shape change vectors is low (*ds*-*neur*: r = 0.11, Figure 2).

**Figure 2.**
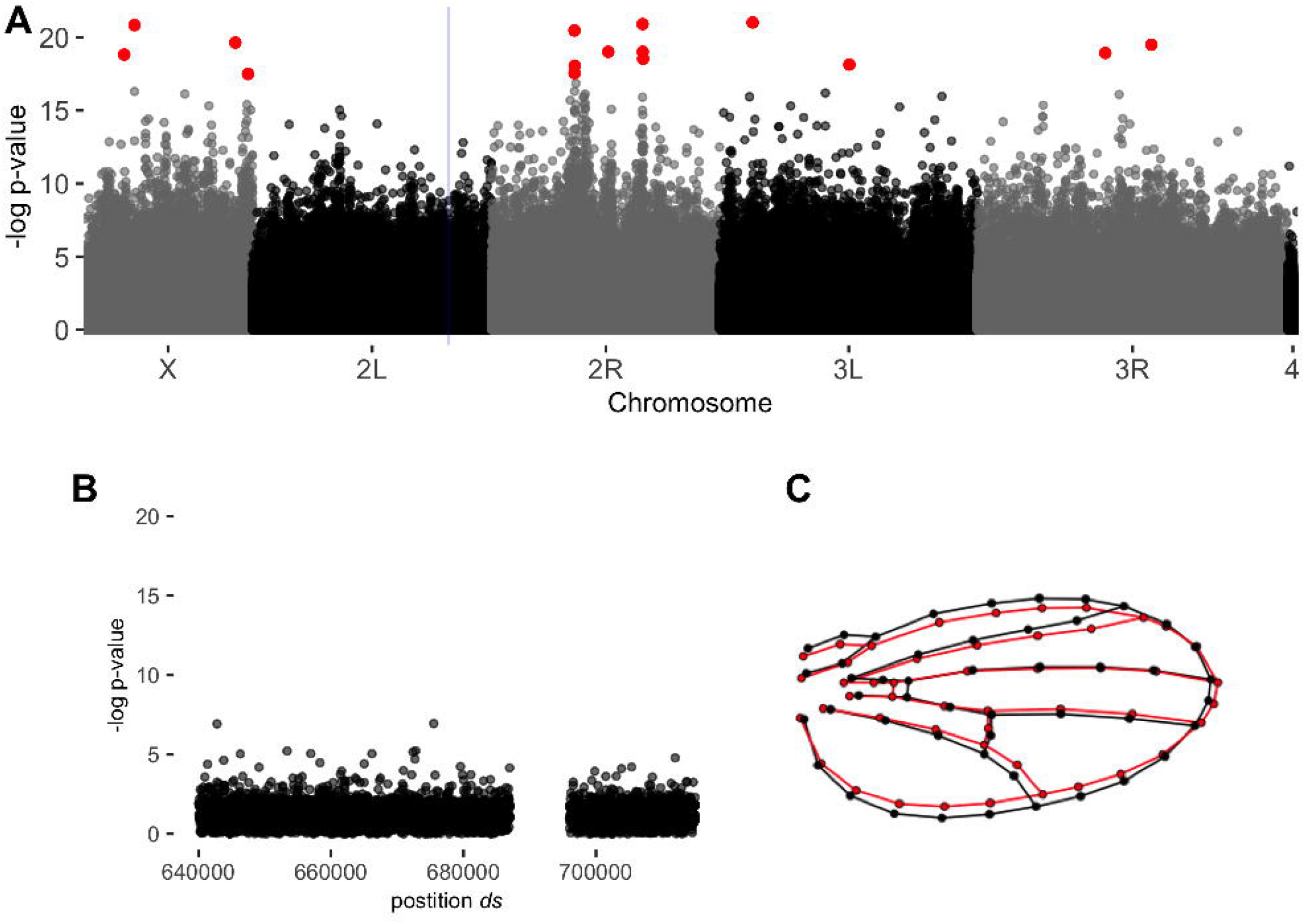
Projections of data onto RNAi shape change vectors are correlated with major axes of shape variation in wild-caught *Drosophila*. Correlations between projection of shape data from CMO population onto *ds, emc* and *neur* RNAi shape change vectors, and the first three eigenvectors from the PCA, calculated from shape data from all samples in the CMO population. PCs 1−3 account for 24%, 18% and 9% of overall shape variance in the CMO population.

### Multiple loci linked to hippo signaling - including *ds*- respond to artificial selection for *ds* and *emc* shape changes

To examine if variants in *ds* are contributing to shape variation, and independently replicate the findings of the earlier GWAS (Pitchers et al. 2019), we performed artificial selection experiment for wing shape along the *ds* shape change vector, and examined the genomic response to selection. By the final generation of selection, we observed a substantial shape change in both the “up” (females: Procrustes Distance (PD) = 0.039, males: 0.044) and “down” directions (females: PD = 0.022, males: PD = 0.022), compared to the base population at the start of the experiment. In comparison, the shape change among unselected control lineages was much smaller (females: PD = 0.005, males: PD = 0.005) (Figure 3, Supplemental Figure 10). The direction of phenotypic shape change after seven generations of selection was in a similar direction to the *ds* shape change vector (defined by RNAi knockdown) for both the up (females: r = 0.90, males: r = 0.90) and down (females: r = −0.82, males: r = −0.77) selection lineages. Realized heritabilities, averaged over sex and replicate were moderate (Supplemental Figure 11, up = 0.38, 95% CI: 0.25 – 0.50; down = 0.28, 95% CI: 0.24 – 0.50). Hippo signaling, including the effects of *ds*, is often associated with changes in size (Pan, 2007). However, we do not observe a significant change of wing size in our selection lineages in either sex (Supplemental Figure 12). It is possible that with more generations of selection we would have observed a clear change in size, as there is a trend indicating such divergence (Supplemental Figure 12).

**Figure 3.**
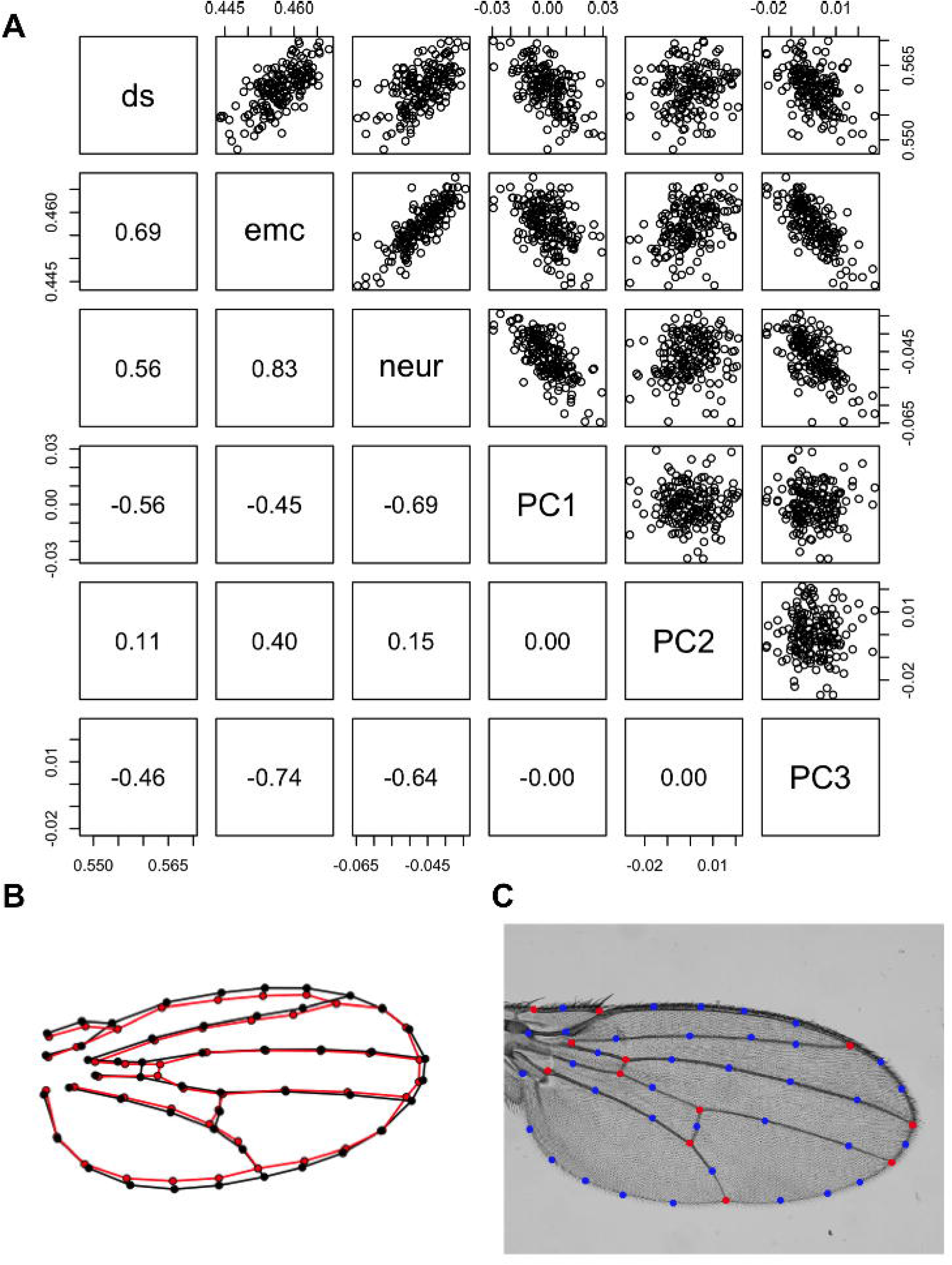
Artificial selection along ds shape change vector influences allele frequencies of variants at *ds.* (A) Phenotypic response to selection based on *ds* shape change vector. Only data from females is plotted for ease of visualization. Each replicate of up (squares), control (dots) and down (triangles) selection lineages are plotted (greys). Estimated response to selection shown along red lines. Wing plots represent the effect of selection on shape change between generation one and seven (red, effects not magnified). (B) Genomic differentiation (F_ST_) between up and down selection treatments measured in 5000bp windows. Red line represents the location of the *ds* locus. Grey line represents 3 standard deviations from genome wide mean F_ST_.

Genome-wide patterns of F_ST_ were examined between up and down *ds* selection lineages. We observed strong genetic differentiation linked with the *ds* locus (Figure 3, Supplemental Figure 13), along with several other regions in the genome. One of the SNPs in the intron of *ds* (2L:702560), identified in Pitchers et al. (2019) through GWAS, showed the expected pattern of response to selection, with opposing sign in up and down selection lineages, with the SNP going to high frequency in all three up selection lineages (Table 1). It should be noted that this SNP is near a complex polymorphism including an insertion of 18bp that may result in inaccurate genotyping at this locus (Supplemental Figure 14). Gene ontology analysis for genes in regions of the genome with an F_ST_ greater than 0. 345 (three standard deviations from mean F_ST_), show enrichment for hippo signaling loci (Supplemental Table 3). The top 20 enriched terms are all related to cell signaling and development. Of note is the inclusion of the terms for ‘negative regulation of hippo signaling’ (GO:0035331), and ‘hippo signaling’ (GO:0035329) in this list (Supplemental Table 3, Supplemental Figure 13). Using a permutation test we confirmed these results, selecting random sets of genomic intervals equal in size to the number of observed outlier windows, and measured the ratio of genes annotated to the expected number of genes in these regions. The observed value for the terms for hippo signaling (ratio = 4.76) and negative regulation of hippo signaling (ratio = 9.23) were in the upper 99.5% percentile in comparison to the distributions under permutation (Supplemental Figure 15).

For the artificial selection experiment based on the *emc* shape change vector we observed phenotypic differentiation under artificial selection in both up (females: PD = 0.043, males: PD = 0.040), and down directions (females: PD = 0.021, males: PD = 0.020), with little change in control lineages (females: PD = 0.009, males: PD = 0.008) (Figure 4). The direction of phenotypic change is correlated with the *emc* (RNAi knockdown) shape change vector in both up (females: r = 0.75, males: r = 0.69) and down (females: r = −0.69, males: r = −0.75) directions. Realized heritabilities, averaged over sex and replicate were calculated for both up and down lineages (Supplemental Figure 16, up = 0.38, 95% CI: 0.29 – 0.47; down = 0.28, 95% CI: 0.21 – 0.35). Genetic differentiation linked to the *emc locus* was modest following selection, but we again observed striking genetic differentiation linked to *ds* (Figure 4, Supplemental Figure 13). Notably, as seen in Supplemental Figure 1, the site frequency spectrum (SFS) suggests modest allelic variation at the *emc* locus in the synthetic outbred population. Using a three standard deviation cut-off for F_ST_, we did observe enrichment for various developmental GO terms, but not of hippo signaling terms (Supplemental Table 4, Supplemental Figure 13).

**Figure 4.**
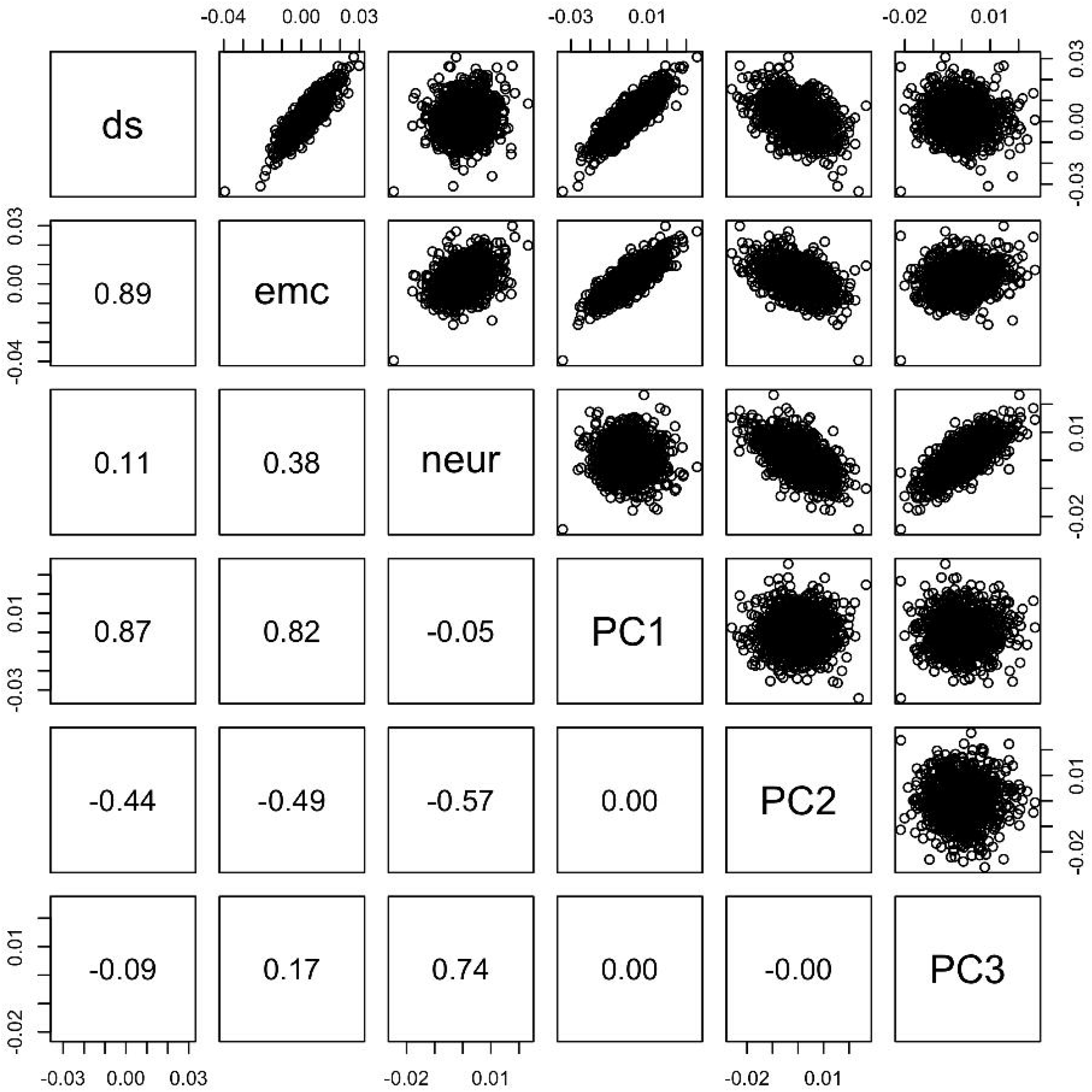
Artificial selection along *emc* shape change vector has modest influence on allele frequencies at *emc*, but a greater impact at the *ds locus*. (A) Phenotypic response to selection based on the *emc* shape change vector. Only data from females is plotted for ease of visualization. Each replicate of up (squares), control (dots) and down (triangles) selection lineages are plotted in greys. Estimated response to selection shown along red lines. Shape change between generation 1 and 7 is indicated on the right. Shape effects have been magnified 5x. (B) Genomic differentiation between up and down selection lineages (F_ST_) measured in 5000bp sliding windows. Red and purple vertical lines represent genomic locations of *ds* and *emc* respectively. Grey line represents 3 standard deviations from genome wide mean F_ST_.

### Bulk segregant analysis in wild caught cohorts does not recapitulate effects of the GWAS or artificial selection

Having demonstrated that variants in (or linked to) *ds* respond to artificial selection for wing shape along the *ds* shape change vector, we next wanted to determine whether we could recapitulate these findings with wild caught individuals. In addition to determining whether we can replicate effects in wild cohorts, it provides the opportunity to identify causal SNPs because of low LD generally observed in wild caught *Drosophila*. Wild caught populations introduce considerably more environmental variation for shape along with a different site frequency spectrum for variants contributing to shape variation (and *ds* like shape changes specifically). In particular, it is known that several of the variants that the original GWAS detected in*ds* have low minor allele frequency (MAF) (Pitchers et al., 2019) (Table 2). The SNP at 2L:702560 does appear to be at intermediate frequency but it occurs both directly before and after an indel, making alignment and variant calling in this region challenging (Supplementary Figure 14). We have included the frequencies (Table 2), but these results should be interpreted with caution due to the technical complexities of mapping and variant calling close to indels.

**Table 2.** *ds* Variants from Pitchers et al. (2019) in wild-caught cohorts used in the present study. Estimated effect sizes for SNPs are estimated from the DGRP GWAS with LASSO regularized coefficients. MAF in wild cohorts was estimated from sequenced pools of 75 random individuals.

| Variant | Estimated Effect (Pitchers 2019) | DGRP MAF | Estimated MAF CMO | Estimated MAF FVW13 | Estimated MAF FVW14 | Estimated MAF PHO |
| --- | --- | --- | --- | --- | --- | --- |
| 2L:655894 | 0.072 | 0.445 | 0 | 0 | 0 | 0 |
| 2L:702560* | 0.159 | 0.056 | 0.375 | 0.473 | 0.485 | 0.336 |
| 2L:702798 | 0.101 | 0.089 | 0.077 | 0.101 | 0.044 | 0.034 |
| 2L:718623 | 0.225 | 0.033 | 0.051 | 0.021 | 0.044 | 0.100 |
| 2L:718627 | 0.11 | 0.033 | 0.055 | 0.020 | 0.046 | 0.099 |

\* This is a complex polymorphism with linked SNPs and INDELs, in Pitchers *et al* (2019) a SNP in this region was found to be linked. However, the variant calling pipeline used in this work recognized an INDEL in this region which was used for counting.

As we sampled multiple cohorts of wild-caught flies in different locations and years in Michigan (USA), we wanted to confirm that any phenotypic differentiation among these samples was modest and would not impact genomic analysis for the BSA. We observe modest, statistically significant wing shape differences among cohorts from a Procrustes ANOVA, utilizing permutations of the residuals for the relevant “null” model (Supplemental Table 5; R^2^= 0.16, F = 351, Z_RRPP_ = 18.3, p = 0.001) (Collyer and Adams, 2018). This appears to be due to differences in wing shape between the PHO population and other populations based on pairwise Procrustes Distances (Supplemental Table 5, Supplemental Figure 3). In a joint PCA including all populations, there is very modest separation between populations using allometry adjusted shape (Supplemental Figure 3). Most relevant to the BSA approach we used, when we project all wild caught individuals onto the *ds* and *neur* vectors, there is no clear separation among sampling locales (Supplemental Figure 7). There is some variation in wing size between populations (Supplemental Figure 8), but this is unlikely to influence downstream analysis as we use size adjusted estimates. There is little evidence of genetic differentiation between populations with the two collections from Fenn Valley Winery separating more on a Principal Co-ordinate Analysis (PCoA)(Supplemental Figure 17) than other sampling locales. There is also no relationship between genetic and phenotypic distances between samples (Supplemental Figure 18). These results suggests that the multiple sampling locales should not influence downstream genomic analysis as individuals used for generating pools were compared within each population, and we observe little evidence for substantial differences among populations.

Because there is a single bout of phenotypic selection distinguishing pools for the BSA, changes in shape and allele frequencies are expected to be modest. We observe shape differences between the two pools within each population (PD = CMO: 0.033; PHO: 0.036; FVW13: 0.040; FVW14: 0.041; Supplemental Figure 19). Correlations of the shape difference vectors of the pools (i.e. difference between the two pools created from the extremes along the *ds* shape change axis), and the direction of the *ds* shape change vector used for selection, is high (CMO: 0.94, PHO: 0.79, FVW13: 0.92, FVW14: 0.90).

BSA genome scans show little evidence of genetic differentiation linked to the*ds gene* (Figure 5). Across the genome, 15 sites were detected as significantly differentiated between “up” and “down” selected pools based on a CMH test with FDR cut-off of 5% (Figure 5, Table 3). The genes nearest to these sites are not associated with hippo signaling pathways or implicated in the development of the *Drosophila* wing (Table 3). Because PHO had somewhat distinct shape variation from the other populations and had a lower correlation of the difference vector between selected pools and *ds* shape change vector, we repeated the CMH test with this population left out. We observe significant differentiation at 174 sites between “up” and “down” pools (Supplemental Table 6, Supplemental Figure 20). We identified the nearest genes to these sites and GO analysis indicated enrichment for wing development terms, in particular related to Wnt signaling, but not hippo signaling terms (Supplemental Table 6). Importantly, we do not observe differentiation linked to *ds* or any other hippo loci. To ensure that the results we obtained were not due to uneven coverage between samples, we down-sampled genomic coverage to 75x for each sample, dropping sites that did not meet this threshold. Significant differences were detected at 19 sites (Supplemental Figure 21, Supplemental Table 7), but none of these overlapped with those identified using all the genomic data. Two of the significant sites are located in the *dumpy* gene, a gene known to have a role in wing morphogenesis during pupation (Etournay et al., 2015). F_ST_ between selected and random pools within each cohort are generally low (Supplemental Figure 22).

**Figure 5.**
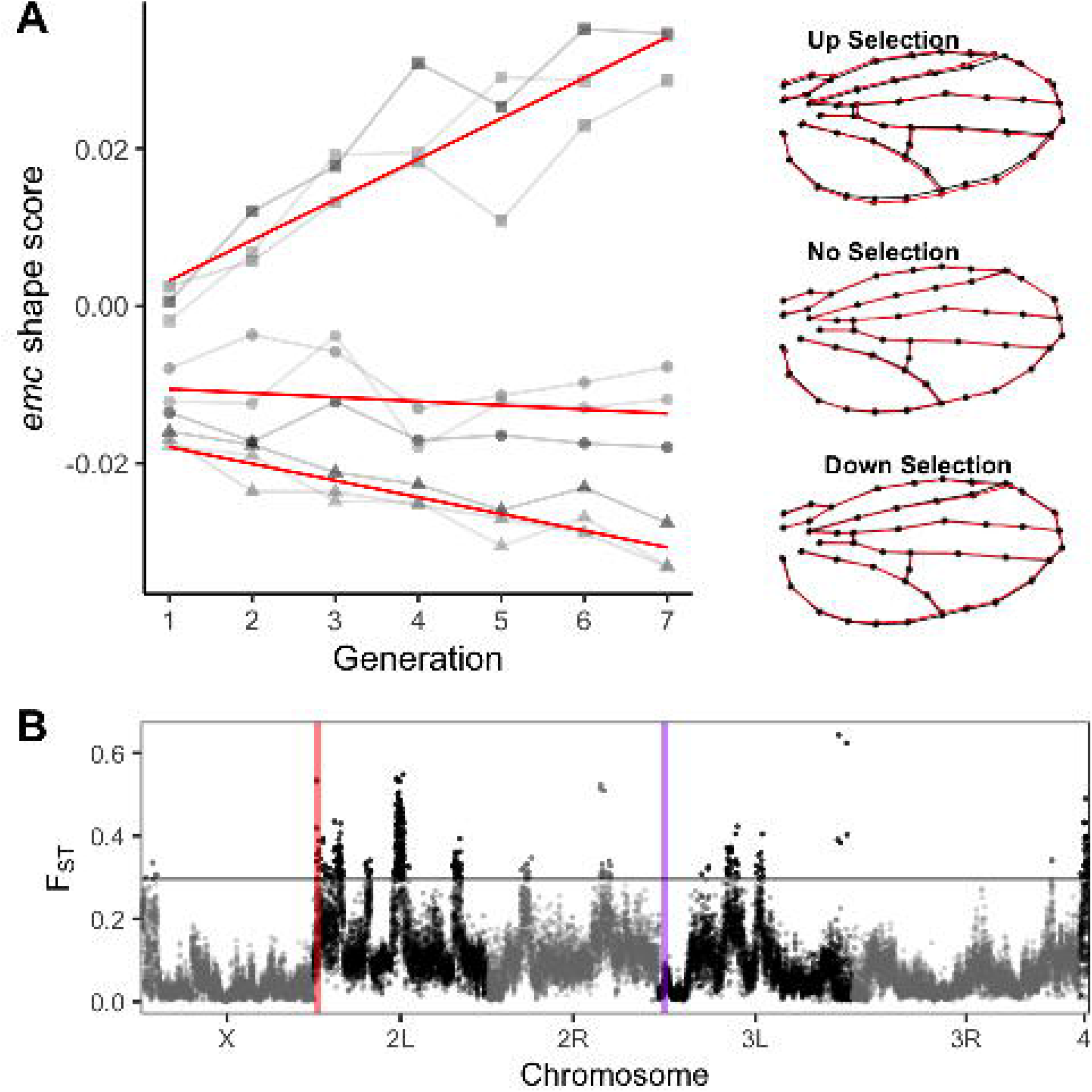
Genetic differentiation between pools selected based on ds shape change among the wild-caught cohorts. (A) Genome-wide scan for differentiated loci between pools selected based on *ds* shape change vector using the CMH test implemented in ACER. Points in red indicate sites with significant differentiation. Position of *ds* gene in blue (B) Genomic differentiation at *ds* between pools selected based on *ds* shape change vector. No sites are significantly differentiated in *ds*. The large gap in sites is due to a masked region in the genome due to repetitive sequence and poor (syntenic) mapping scores. (C) Shape difference between selected pools of individuals from one representative (CMO) population, with the mean shape of pools represented in black and red.

**Table 3.**
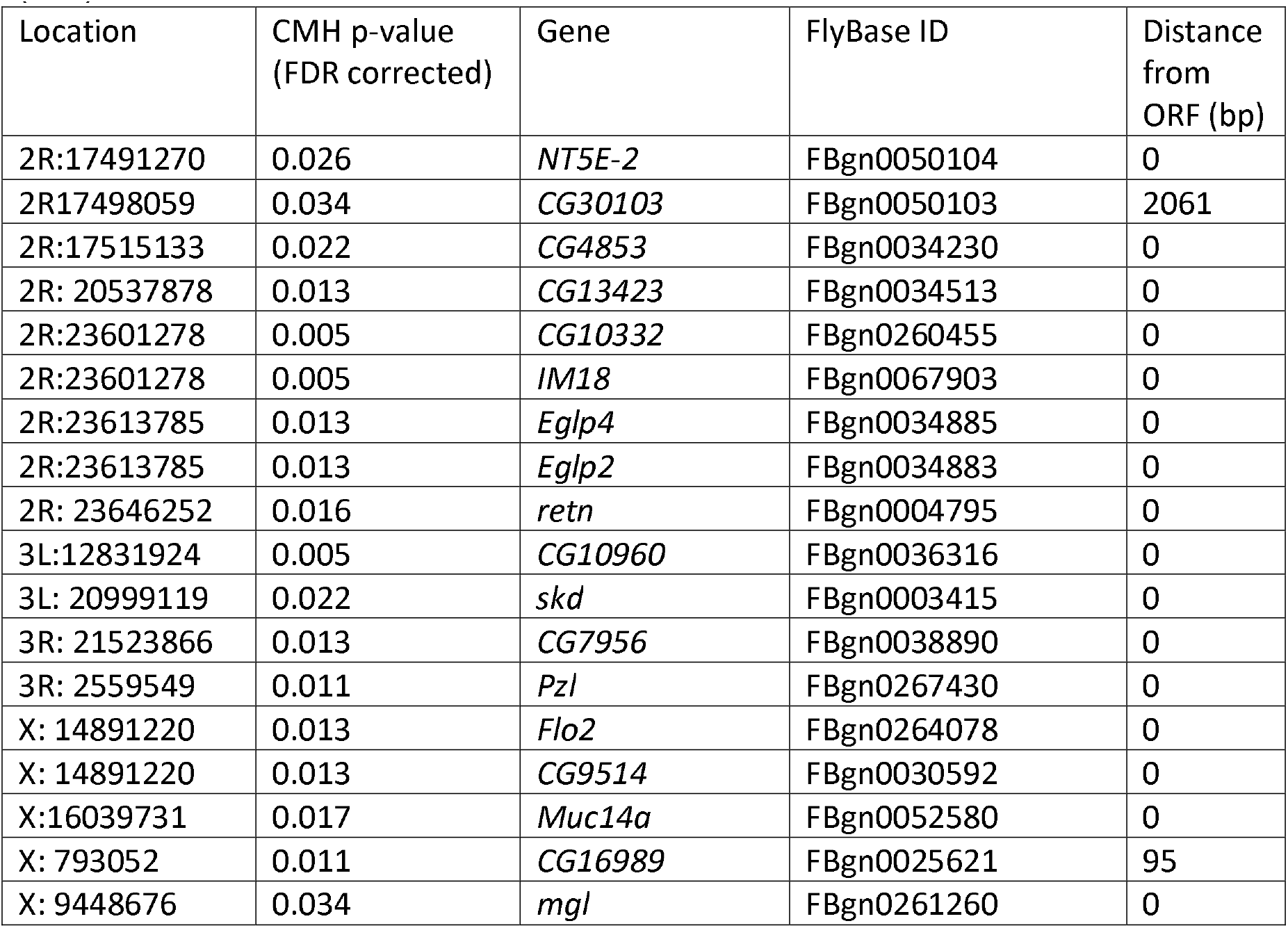
Significantly differentiated variants for *ds* shape change from the wild-caught cohorts (BSA).

In addition to the BSA selection based upon the *ds* shape change, we also selected pools of individuals based on the *neur* shape change vector. We did not use *emc* shape change in this experiment due to the high similarity between the *ds* and *emc* shape change vectors (r = 0.65), and the similar response to selection reported above. We selected the *neur* shape change vector as it is not aligned with *ds*, but does align with directions of natural variation, in wild populations (Figure 1, 2, Supplemental Figure 9). Additionally, there is little relationship between the *ds* and *neur* shape change axis (r = 0.12, Supplemental Figures 7 and 8) in the wild caught cohorts. We observe shape changes between pools of individuals (PD = CMO: 0.027; PHO: 0.028; FVW14: 0.041; FVW13: 0.038, Supplemental Figure 23). There is little evidence of genetic differentiation between *neur* selected pools (Supplemental Figure 24). Only 4 sites were identified as being significantly differentiated between pools and none of these sites are associated with wing development (Supplemental Table 9). When population differentiation between pools within populations is measured using F_ST_, genetic differentiation remains low across the genome (Supplemental Figure 25)

## Discussion

The primary goal of this study was to determine whether we could recapitulate genetic effects initially observed through a traditional GWAS using an “inverted” approach: artificially selecting on phenotypes and observing changes in allele frequencies. We observed that shape changes associated with the *ds*, *emc* and *neur* genes were associated with major axes of genetic variation among a panel of wild type strains (DGRP) reared in the lab, and axes of phenotypic variation among wild caught individuals (Figure 1, 2). After observing a strong response to artificial selection along two shape change vectors (*ds* and *emc*), we examined patterns of genomic differentiation and observed substantial changes in allele frequency for markers linked with *ds* itself (figure 3), and markers linked to numerous genes associated with hippo signaling (Supplemental Figures 13 and 15).

In contrast, our BSA experiments, using pools of wild caught individuals chosen to be phenotypically divergent on the same shape vectors, did not detect differences in the loci identified in the artificial selection experiments (Figure 5, Supplemental Figure 24). As we discuss in detail below, these seemingly contradictory results are in fact not that surprising.

Following artificial selection based on *ds* shape change we observe allele frequency changes not only at *ds* but also linked to a number of other hippo signalling loci (Figure 3, Supplemental Table 3). The previous GWAS study identified a number of loci associated with wing shape variation in the DGRP, however, this approach cannot predict which alleles are causative (Pitchers et al., 2019). In our synthetic outbred population, we maximized variation among haplotype blocks containing many of the candidate SNPs in *ds*, increasing our ability to detect frequency changes at and near the implicated variants. Although LD blocks in the outcrossing population from this study remain large, *ds* variants exist on multiple distinct haplotypes, allowing for an examination of allele frequency changes for each. Of particular interest is SNP 2L:702560, previously identified though GWAS (Pitchers et al., 2019) as influencing wing shape variation. It was driven to near fixation in each of the artificial selection lineages (Table 1). Although this polymorphism is annotated as a SNP, this region may contain a complex polymorphism (Supplemental Figure 14), making it difficult to accurately assess genotypic calls. Because of this, the predicted allele frequency in the founding population and allele frequencies in this region may be inaccurate. Previous studies demonstrate the importance of alleles at intermediate frequency in founding populations to those contributing to responses to selection over short timescales (Kelly and Hughes, 2019). If this polymorphism is at a more intermediate frequency in the founding population, it would be more likely to be captured by selection during these experiments. Additionally, haplotype blocks in the initial population are large, and may contain many potential functional variants. However, based on the results of both the current and previous studies, these *ds* variants associated with 2L:702560 are good candidates for functional validation in future work.

When selecting on the *emc* shape change vector, which is similar to that of the *ds* shape change, we observe only a modest allele frequency change at *emc*, and a more robust response at *ds* (Figure 4). In hindsight, this is not particularly surprising and there are multiple contributing factors. Given the increased genetic diversity at *ds* compared to *emc* in the founding population, alleles in *ds* may have provided a more accessible genetic target, as selection can only act upon the diversity available in the population. Additionally, if our estimated direction of effects and selection for *emc* (based on RNAi knockdown) was not well aligned with the actual direction of *emc* SNP effects, this could result in weaker selection on variants at the *emc* locus. It is worthwhile pointing out the small magnitude of the *emc* shape change vector (0.44) relative to *ds* (5.5). However, previous work has indicated that there is a relationship between this estimated *emc* shape change vector (from RNAi) and the effect of SNPs in *emc* on shape change (Pitchers et al., 2019).

In addition to a response on allele frequency associated with *ds*, our results suggest a response on segregating variation at other hippo signaling loci in the *ds* artificial selection experiment. Earlier work has suggested that the direction of effects within signaling pathways are inconsistent for alleles of small effect (Dworkin and Gibson, 2006). However, allelic effect sizes in the 2006 study were heterogeneous and may result in direction and magnitude being confounded. In contrast, in both the current and the Pitchers et al. (2019) studies, we estimated the direction of genetic effects by titrating gene knockdown. The strength of this approach is highlighted in the result that segregating variation at multiple hippo loci was selected on (Supplemental Figures 13 and 15). Our finding is consistent with models for the architecture of complex traits that predict that many alleles of small effect will contribute to trait variation with many genes within developmental pathways (Boyle et al., 2017; Wray et al., 2018). This pathway response has also been demonstrated in human adaptation to pathogen resistance (Daub et al., 2013) and high altitude (Gouy et al., 2017). These results are consistent with the expectation that polymorphisms in the same developmental pathway would show correlated phenotypic effects and therefore correlated genomic responses to selection. However, this may not be reflective of all wild caught populations. In this study, we generated a population that had high diversity at *ds*, while these variants are at much lower frequency in natural populations (Table 2). The amount of selectable variation a variant provides, depends on both effect size, *a*, and variant frequencies, *p*, as *V*_A_*= 2p(1-p)a*^2^. When allele frequencies are near 0 or 1, even variants with large effects will have only a small contribution to short term selection response. Therefore, the outcrossed population we created here is an ideal situation to validate the existence of the measured effects. It is unlikely to be typical of natural populations where functional variants may be rare.

Given the clear and robust response observed in the artificial selection experiment, it may seem surprising that we do not observe allele frequency changes in the BSA using the wild cohorts. Indeed, previous work has demonstrated that variants in *Egfr*, could be replicated in wild caught samples (Dworkin et al., 2005; Palsson et al., 2005) and were also found in genome wide associations (Pitchers et al., 2019). However, there are many explanations for why we may not have been able to detect these allele frequency changes in our experiment. First, the addition of environmental variation to the system introduces additional complications. In the aforementioned example with *Egfr*, the genetic effect of the SNP in wild-caught cohorts was ∼10% of the magnitude estimated in lab-reared flies. As discussed previously, the*ds* variants implicated in the previous GWAS study are at low frequency in the natural cohorts (Table 2). Given that natural populations of *Drosophila* are generally large and wing shape is likely under weak selection (Gilchrist and Partridge, 2001), mutation-drift-selection balance may maintain most variation, resulting in low minor allele frequencies at these sites. Because allelic contribution to wing shape are expected to be both rare in wild populations and of small phenotypic effect, we do not expect large allele frequency changes given only one “generation” of selection. Using the approach of ACER (Spitzer et al., 2020) to account for sampling effects, we observe few differentiated sites, and none in the *ds* gene, indicating that BSA may not be well-suited to identify modest allele frequency changes, thus, not particularly effective for polygenic traits. Although our approach was tailored to look for variants that had consistent direction of frequency changes across the four collection cohorts, it is possible that different loci were contributing variation within each cohort. We attempted to address this question by examining allele frequency changes between selected pools within each cohort (Supplemental Figures 22 and 24) but could not identify specific loci contributing to differences within any one population. Previous successful BSA studies identified smaller numbers of contributing loci with few polymorphisms contributing to the trait of interest. For example, in *Drosophila,* a number of melanin synthesis genes contributing to variance in pigmentation between populations were identified using a BSA (Bastide et al., 2013). Pigmentation may represent a relatively ‘simpler’ genetic architecture (fewer variants of individually larger genetic effect, smaller impact of environmental variation, smaller mutational target size) and if so, this may have enabled the success of the BSA approach with such systems. In the case of wing shape, we know that many alleles of small effect contribute to variation in the trait (Pitchers et al., 2019).

Our approach for the BSA was to perform the same phenotypic selection within each of four distinct “populations”. It is important to recognize that there was heterogeneity among our populations, not only in allele frequencies, but in environmental variance and potentially *G*x*E*, even though all were caught in locales in lower Michigan. We detected small degrees of phenotypic and genetic differences between cohorts, however these effects are neither correlated with one another, nor related to the *ds* and *neur* shape scores used for selecting individuals (Supplemental Figures 3, 7, 18). The population from the Phillips Orchard (PHO) was phenotypically distinct from the other populations. When we performed the BSA without this population, we observed a larger set of variants associated with shape (Supplemental Figure 20), albeit still not showing any effects at *ds* or *neur* genes themselves. One possibility is that the increased number of sites when the PHO sample is removed from analysis represents an unknown statistical artefact we have not identified. However, a more likely explanation is that there are some large unknown environmental influences (*E*), or that the genetic effects show a degree of *G*x*E* (with a specific environment in PHO) that contributed to shape variation along the *ds* direction in this population. Such obfuscating effects have been observed before with the previously discussed *Egfr* example, where the SNP effect identified and validated in multiple contexts (Dworkin et al., 2005; Palsson et al., 2005; Palsson and Gibson, 2004) could not be detected in one natural population, despite being at intermediate frequencies in each sample (Palsson et al., 2005). Importantly, we did detect differentiation at sites associated with developmental processes in the wild cohorts, suggesting that the failure to detect variation linked to *ds* or other hippo signaling loci (Table 3, Supplemental Table 6,7) is not due simply to a lack of power.

The response to selection at *ds* and other hippo signaling loci in the artificial selection experiment based on *ds* shape change indicates that this is an important axis of variation for wing shape. Coupled with the alignment of phenotypic effects of perturbations in genes in this pathway with directions of G and P, this finding may seem to suggest a developmental bias in available variation. However, we caution against such interpretations based solely on the findings in this study. The structure of the G matrix strongly influenced our findings as we artificially created a population to maximize genetic diversity at *ds*. When another effect is aligned with *ds* shape change, as in the case of *emc* shape change, we observed the same response at the hippo signaling loci and not at *emc.* Only the genetic diversity in the starting population was available to be selected on so this influenced selection towards the “spiked in” *ds* variants, even if the inferred phenotypic effects of *emc* variants are very similar. Alternatively, the inferred *emc* direction of effects (via RNAi knockdown) may be sufficiently “distant” from true effects of *emc* variants. If this was the case, we were ineffectively selecting for *emc* shape changes. In other cases where single genes are implicated in divergence between multiple populations, such as *mc1r* in mice (Steiner et al., 2007) or *pitx1* in stickleback (Chan et al., 2010), other factors such as low pleiotropy, developmental and mutational constraints and history of selection in the population are used to explain why these genes are so often implicated in evolutionary change (Gompel and Prud’homme, 2009; Martin and Orgogozo, 2013; Stern and Orgogozo, 2008). In our case, it is not *ds* itself that is special but rather the orientation of the G matrix to align g_max_ with the direction of effect for *ds* that shapes our results. Selection acts on variants aligned with the vector of selection (Reddiex and Chenoweth, 2021). By varying the orientation of g_max_ in the parental population, we would be able to address questions about the repeatability of hippo overrepresentation and if this can be explained by more than just the orientation of G.

Despite the need for skepticism about the potential for developmental bias influencing directions of variation, the correlated response of sites linked to multiple other hippo signaling genes is intriguing. Coupling of more traditional mapping approaches like GWAS with short term artificial selection provides an additional route to validation and replication of genetic effects. It also suggests that using multivariate data to address the distribution of genetic effects will pay long-term dividends to our understanding of both inheritance and the evolution of multivariate traits.

## Supporting information

SuppTable3

SuppTable4

SuppTable6

SuppTable7

## Acknowledgements

We would like to thank the Hosken lab for *D. simulans* wing shape data. Yun Bo Xi aided with the sequencing of *ds* haplotype. We thank Dr. Brian Golding for computational resources. We thank Dr. Catherine Peichel and two anonymous reviewers for helpful feedback on the manuscripts.

