## SupplementalFiguresTables for "Complexities of recapitulating polygenic effects in natural populations: replication of genetic effects on wing shape in artificially selected and wild caught populations of *Drosophila melanogaster*"

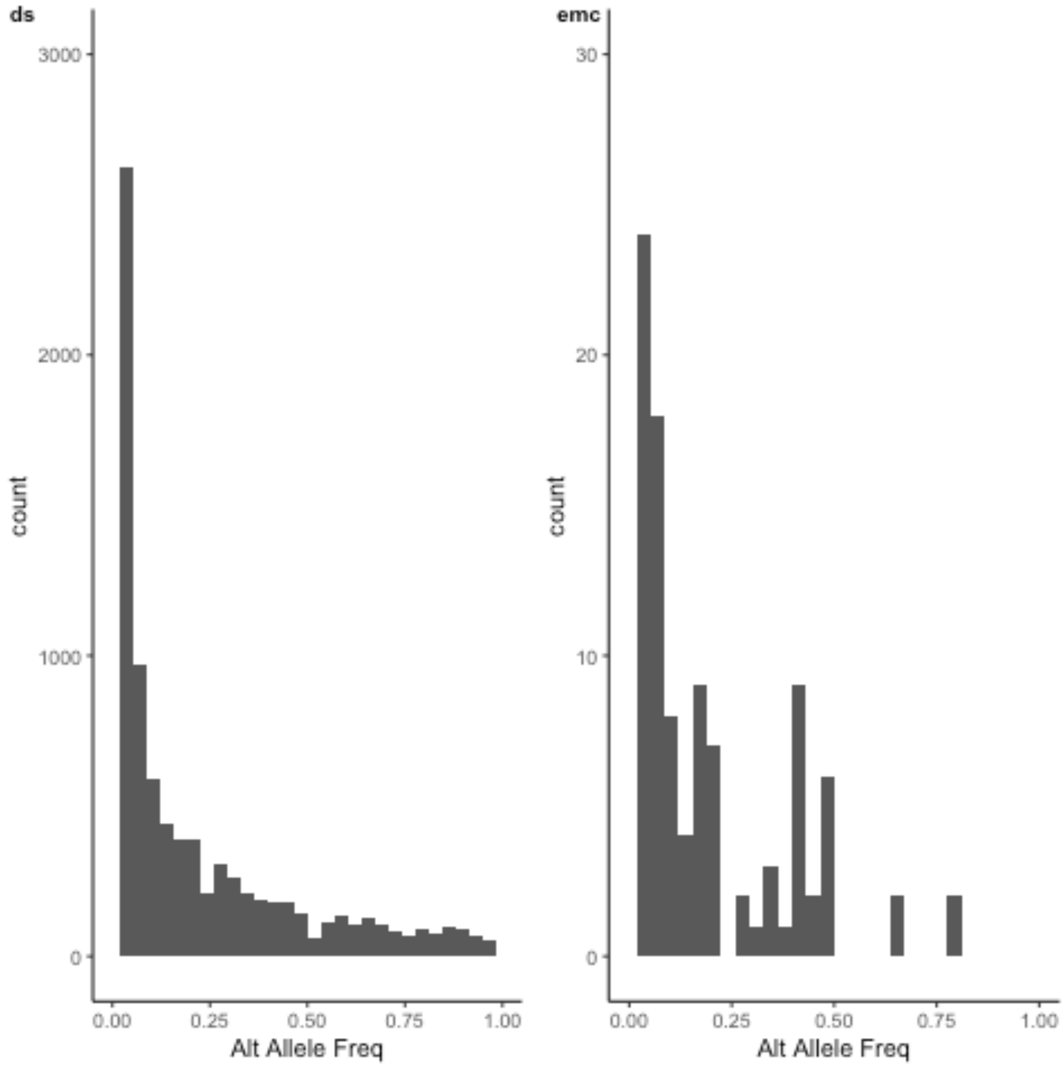


Supplemental Figure 1. Allele Frequency spectra demonstrate more variation at *ds* compared with *emc* in the synthetic outbred population used for artificial selection. Figures show estimated alternate allele frequencies at *ds* and *emc*. Alternate allele frequencies are estimated using parental strain genotype data and assuming an equal contribution from each parent to the founding population. Note the different y axis scale between *ds* and *emc*.

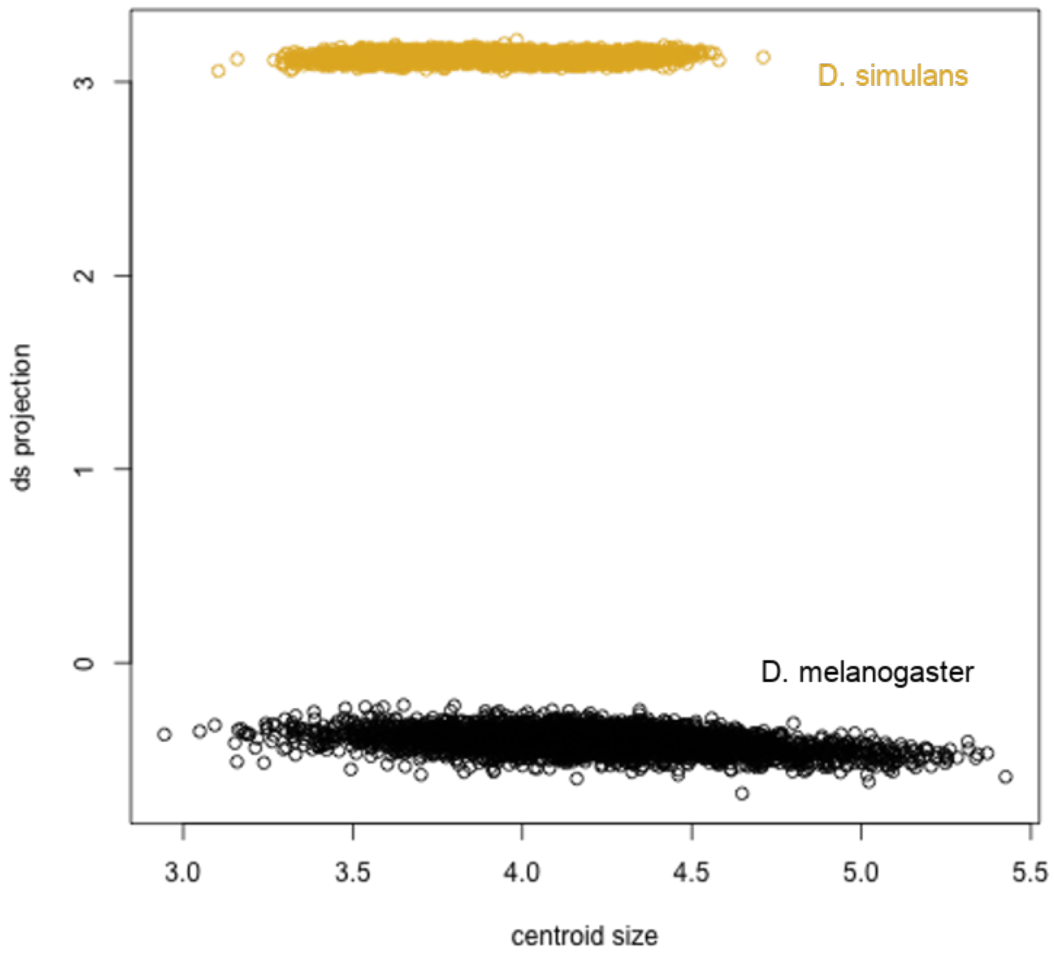


Supplemental Figure 2. Projection of FVW14 wings onto *ds* shape change vector shows clear distinction between *D. melanogaster* and *D. simulans.* Clear separation between the FVW14 samples (black) and *D. simulans* data (gold) indicates that *D. melanogaster* females were accurately identified.

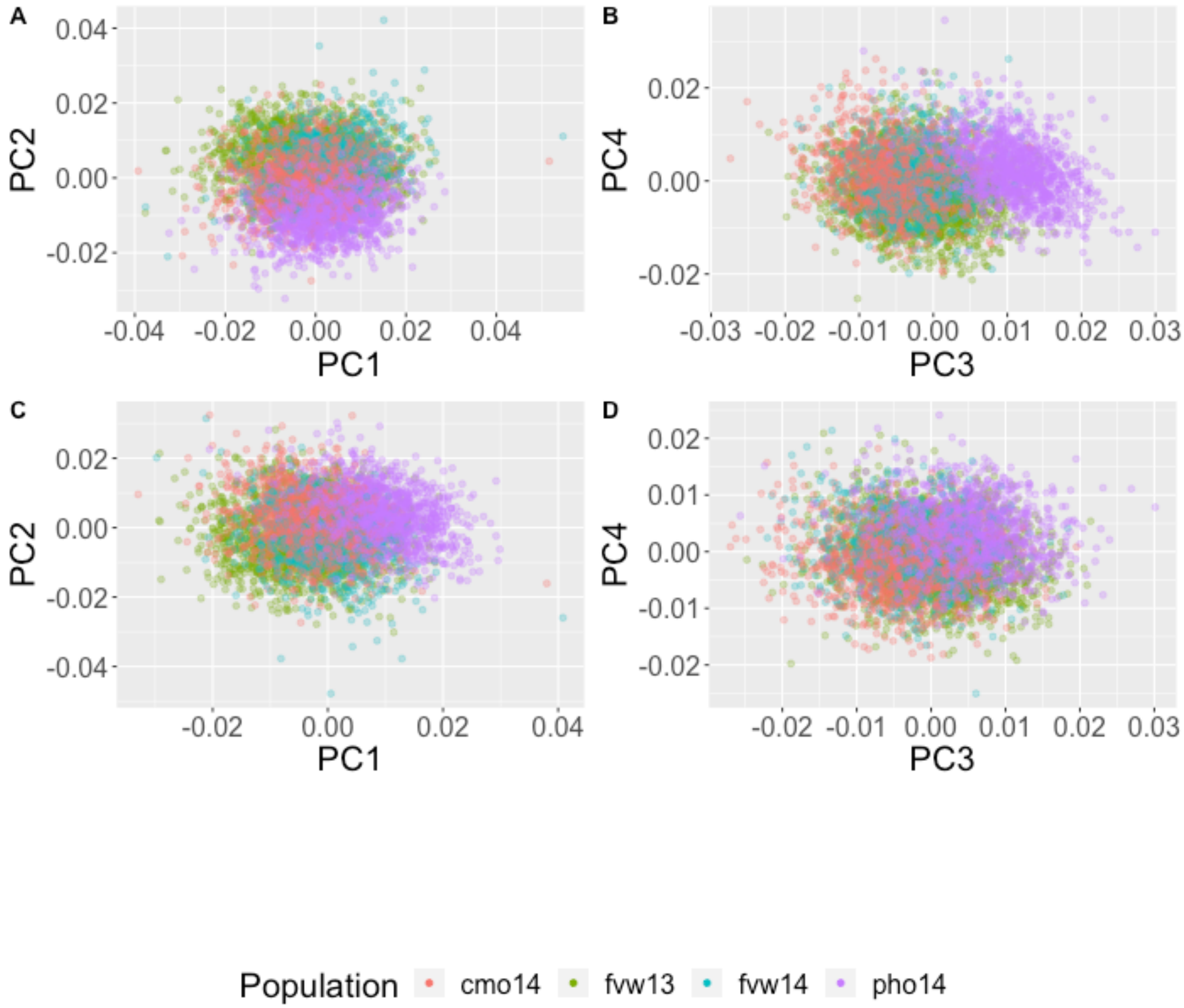


Supplemental Figure 3. Principal component analysis of shape variation within and among populations for wild collected *Drosophila*. First four axes from the PCA for shape variation are shown. (A) and (B) PCA includes all shape variation. (C) and (D) use the 'allometry corrected' landmarks, (residuals from a model regressing landmarks onto centroid size of wings).

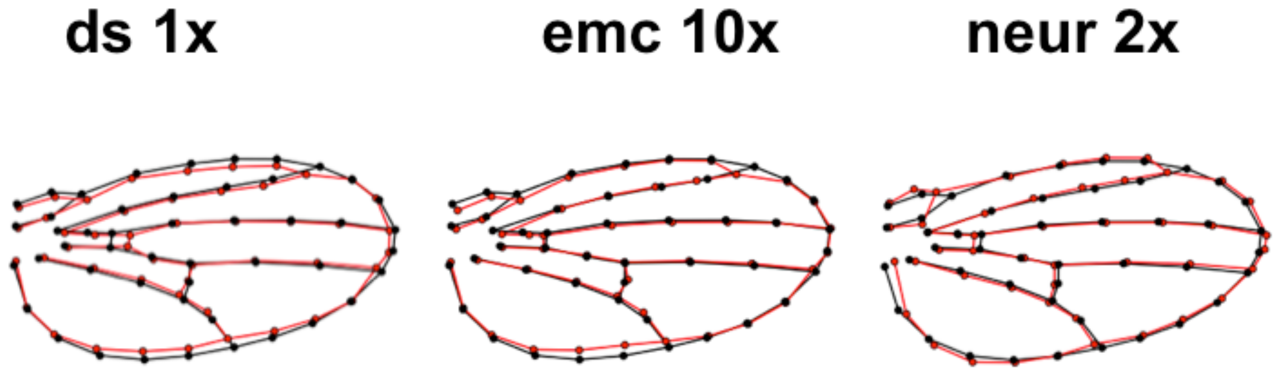


Supplemental Figure 4. Shape change effects due to RNAi knockdown of *ds*, *emc* and *neur.* Scaling of effects provided to aid in visualization of shape changes. Different magnifications are provided to account for the disparate magnitudes for estimated shape change vectors: *ds* = 5.5, *emc* = 0.44, *neur* = 2.8. The vectors from these analyses were used for projections in this study.

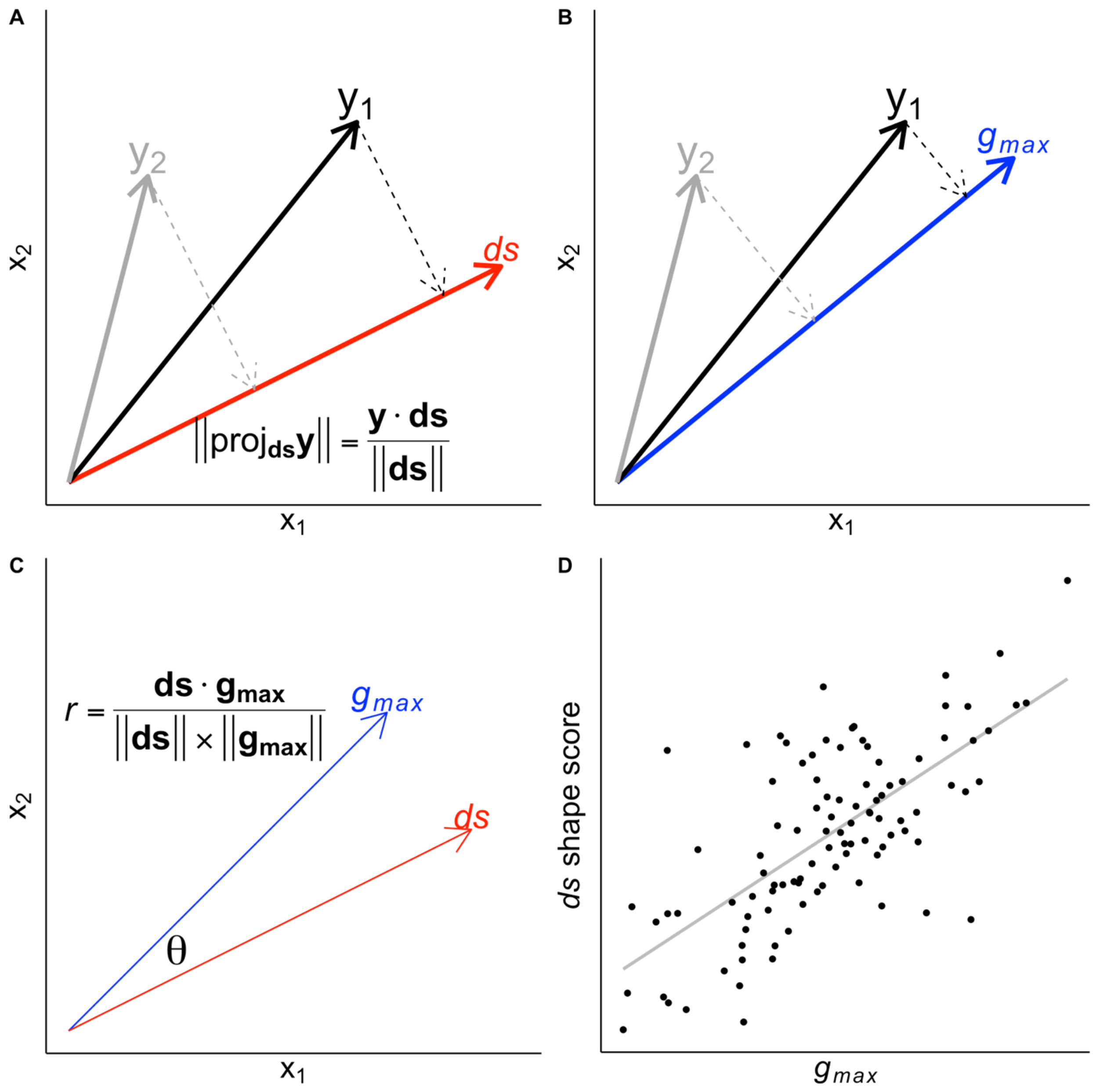


Supplemental Figure 5. Illustration of projections onto shape vectors to generate shape scores used in this study. The *ds* shape change vector is used for demonstration. (A) Calculating shape score using projection. $y_1$ and $y_2$ represent vectors of landmarks for two representative individuals. Dotted arrows represent projection of $y_1$ and $y_2$ onto the shape change vector defined by *ds* RNAi to generate the *ds* shape score. (B) Similarly, $y_1$ and $y_2$ are projected onto the $\boldsymbol{g_{max}}$ (PC1) vector, representing the direction of maximum genetic variation in the genetic variance-covariance matrix, ***G***. (C) Comparing $\boldsymbol{g_{max}}$ and shape change vectors, using the correlation (*r*) of vectors directly, or via the angle θ, between vectors. (D) Hypothetical relationship between *ds* shape scores and $\boldsymbol{g_{max}}$, indicating a relationship between directions of *ds* induced shape change and the direction of genetic variation in shape variation.

A B

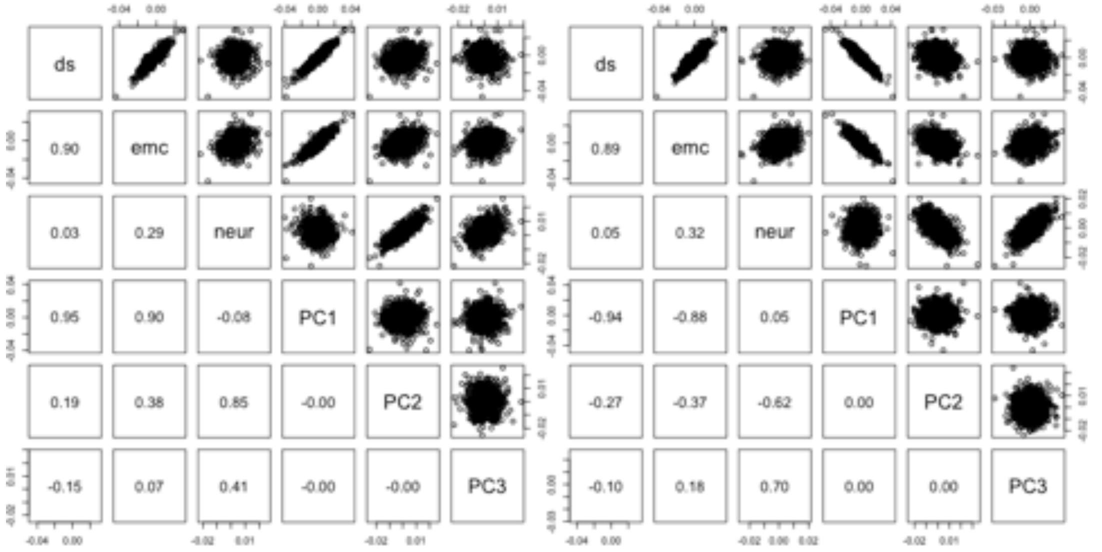


Supplemental Figure 6. Including wild-caught females in the analyses does not change interpretation for FVW14 data. Correlation between projection of shape data from FVW14 population onto *ds, emc* and *neur* shape change vectors and the first three PCs are calculated from shape data from all wings in the FVW14 population. (A) male only data, (B) females and male data. Inclusion of female data does not change the conclusions drawn from the relationships between directions shape change vectors and PCs. Note that the “flipped” directions of the PCs between (A) and (B) represents the arbitrary sign of the set of eigenvectors for the PCA (i.e. the set of eigenvectors can equivalently be multiplied by -1).

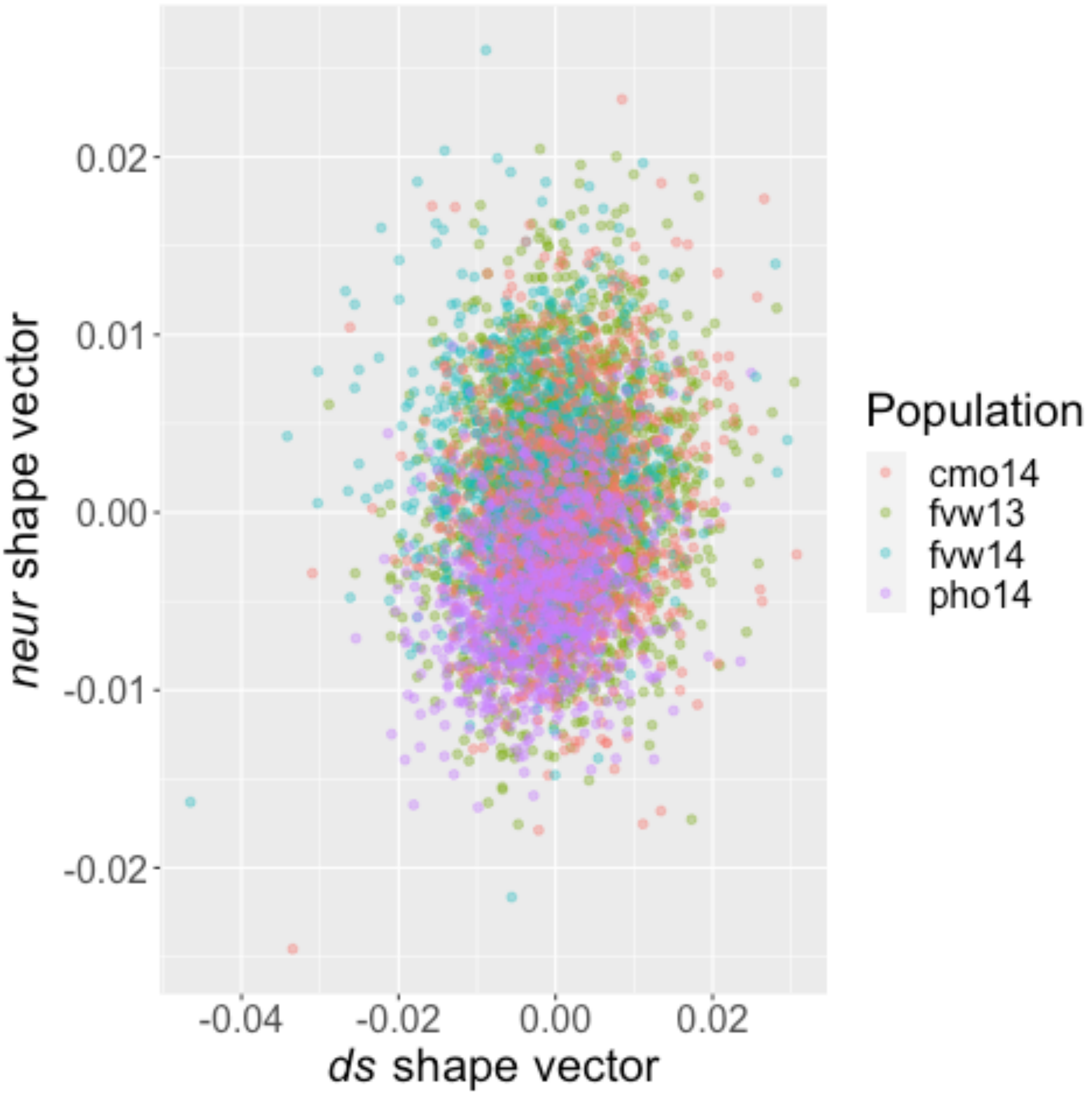


Supplemental Figure 7. Shape variation in field collected samples by projecting individual shape data onto *ds* and *neur* shape change vectors. Projections were performed used size "adjusted" landmark data onto *ds* and *neur* shape change vectors. The correlation between the ds and neur shape scores is r = 0.12. In linear models with shape scores (*ds* and *neur* respectively) regressed onto population and wing size, the partial $R^2$ for population effects is 0.040 (*ds*) and 0.18 (*neur*).

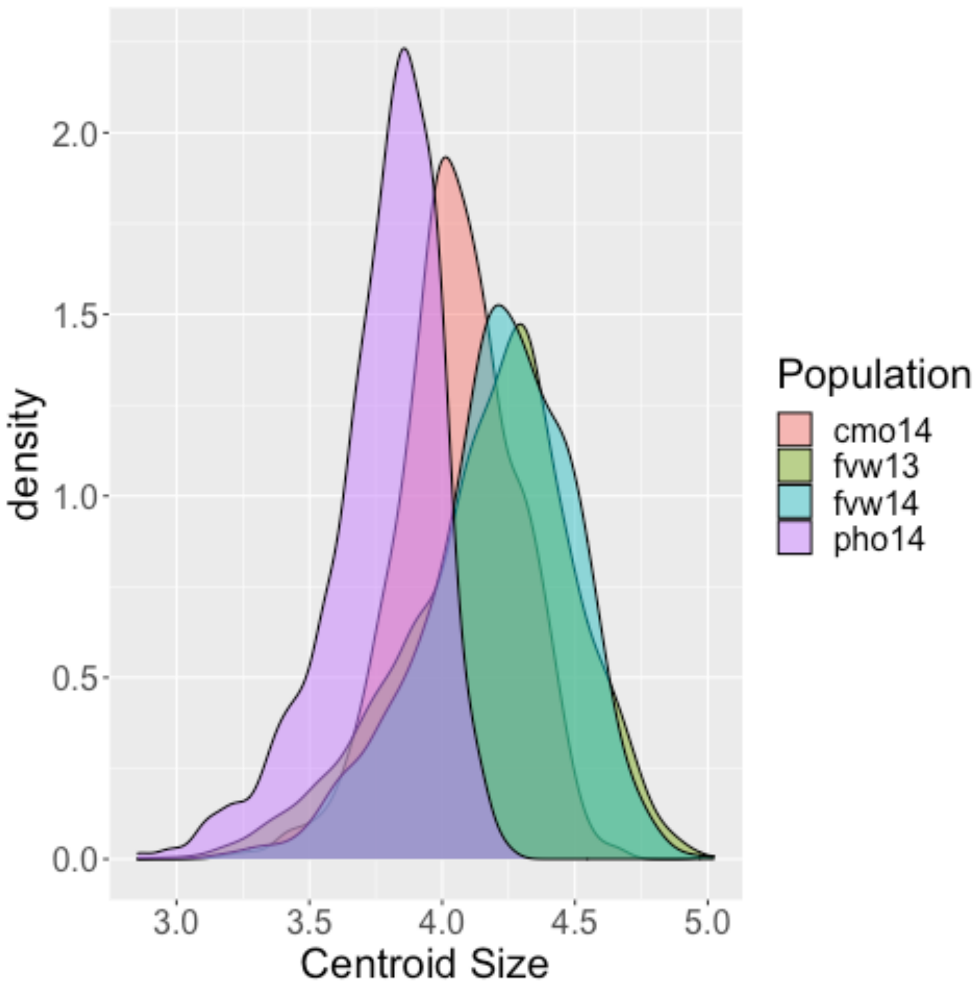


Supplemental Figure 8. Distribution of wing sizes (males only) in wild caught cohorts. When size is linearly regressed onto population, model $R^2 = 0.27$.

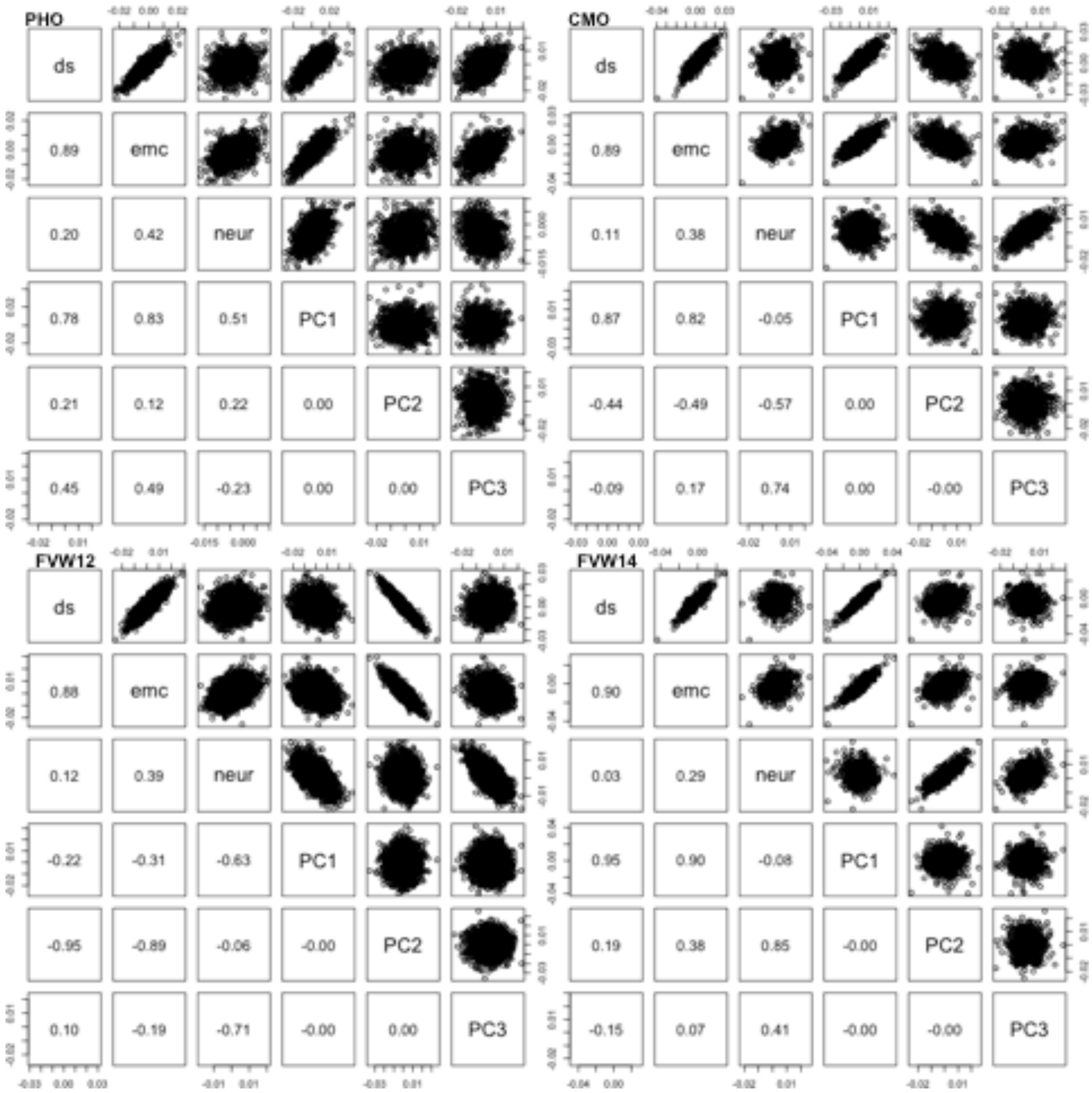


Supplemental Figure 9. **Projections of data onto RNAi shape change vectors are correlated with major axes of shape variation in wild-caught *Drosophila* from each population**. Correlation between projection of shape data from wild cohorts onto *ds, emc* and *neur* shape change vectors and the first three PCs are calculated from shape data from all wings in each cohort independently. Note that the "flipped" directions of the PCs for FVW12 represents the arbitrary sign of the set of eigenvectors for the PCA (i.e. the set of eigenvectors can equivalently be multiplied by -1). Note that the eigenvectors representing PC1 and PC2 in FVW12 have "swapped" because of similarities in variance accounted for.

A B

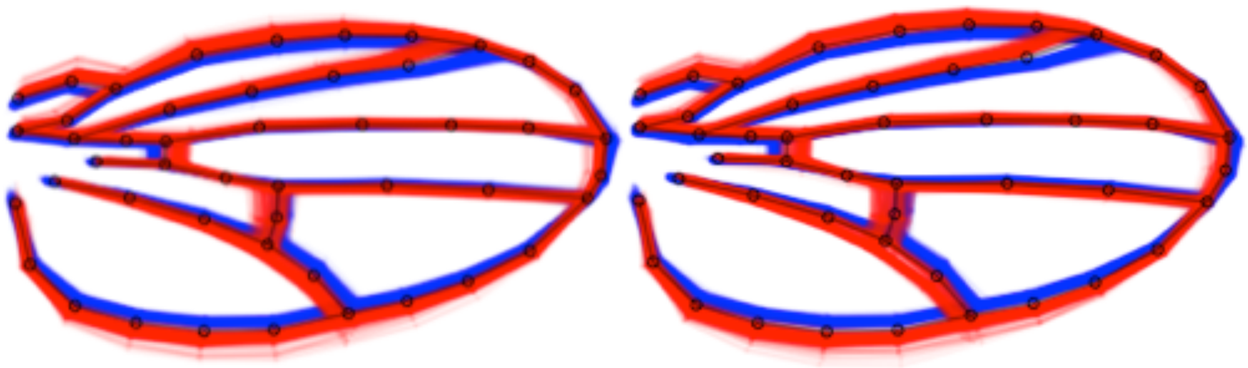

Supplemental Figure 10. Variation in wing shape among individuals from artificial selection along the *ds* shape change vector. All wings from generation 7 of up (red) and down (blue) selection lineages plotted, with females in (A) and males in (B). Black wireframe is mean shape from the experiment.

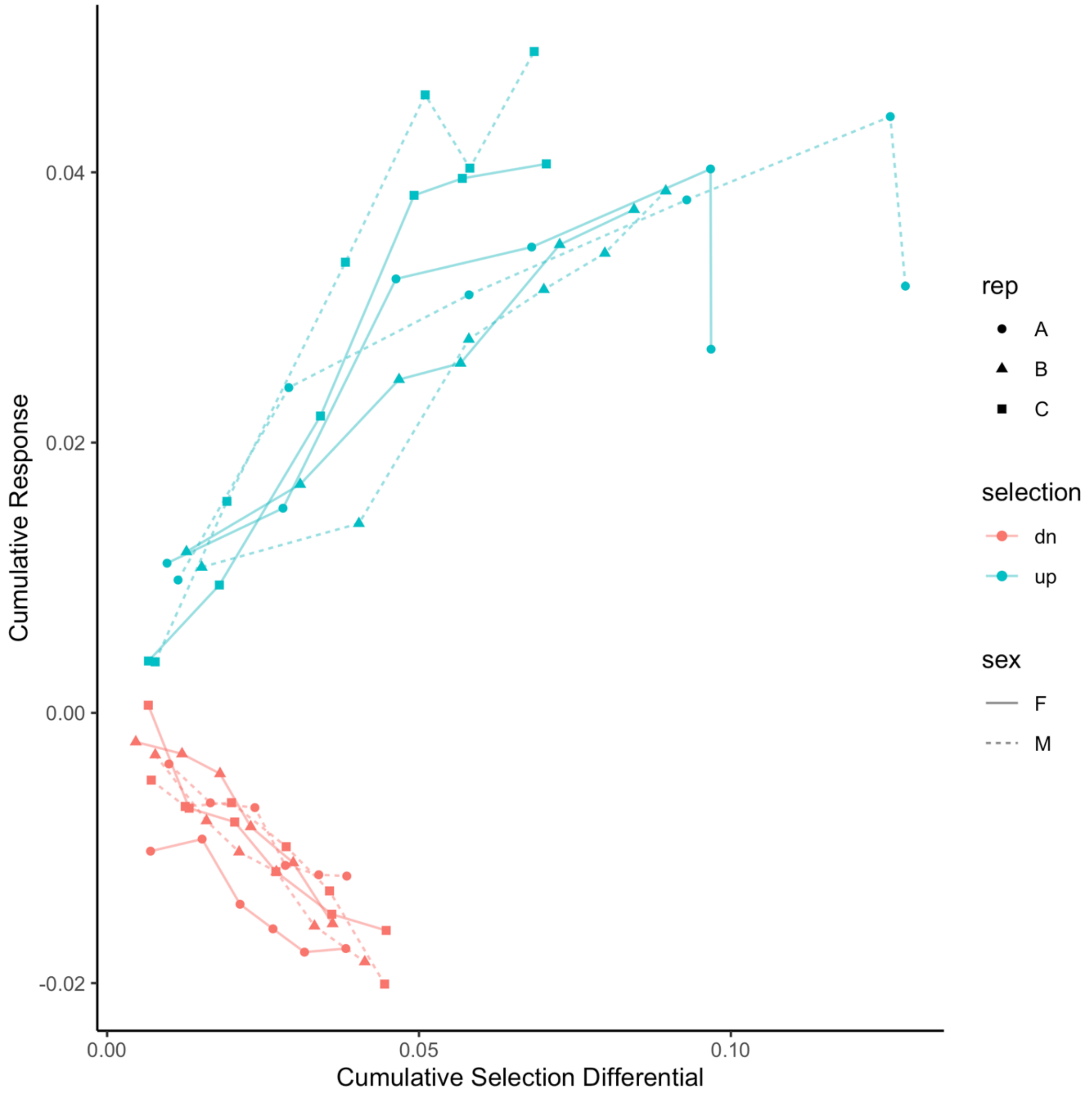


Supplemental Figure 11. Response to artificial selection along the *ds* shape change vector. Regression of cumulative selection differential onto cumulative response was used to estimate realized heritability in both treatments (“up” & “down”) independently.

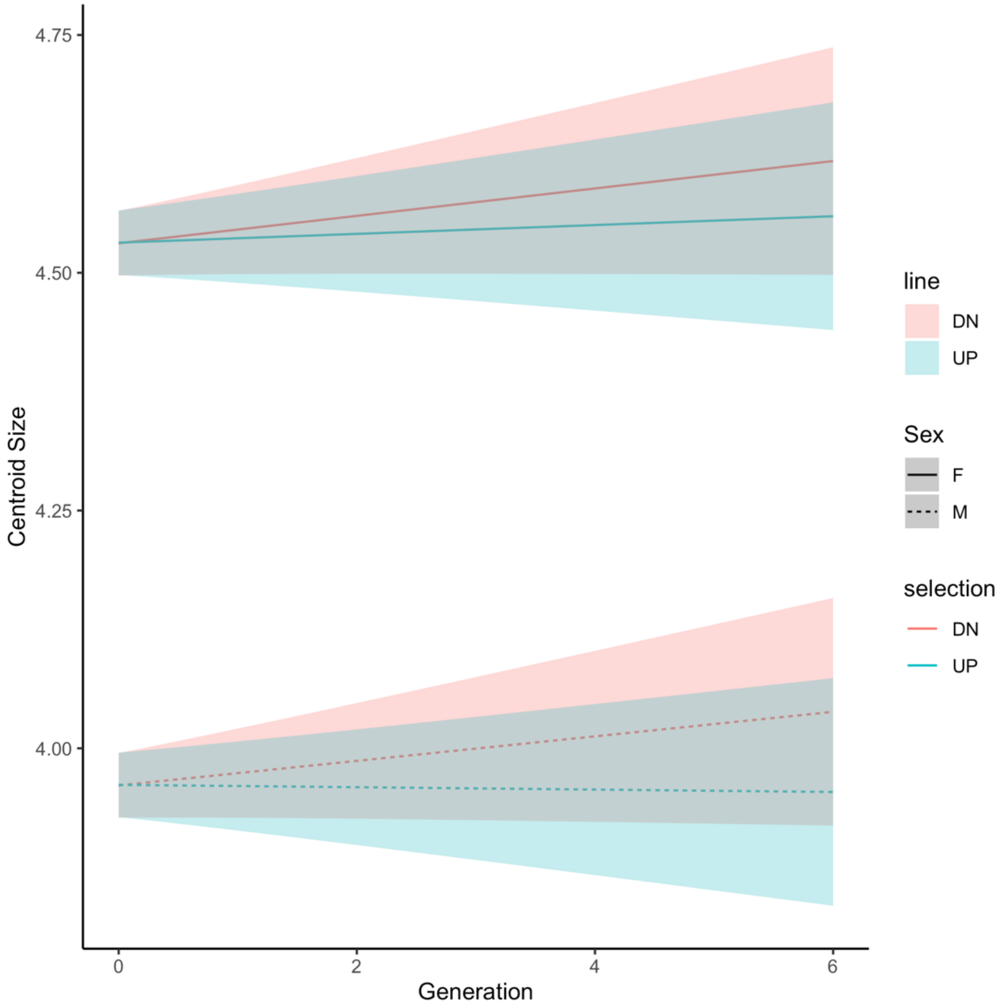


Supplemental Figure 12. Modest (and not significant) changes in wing size following artificial selection based on *ds* shape change. Mean size estimated for up and down *ds* selection lineages estimated from a linear mixed model with replicate lineage fitted as a random effect. Shaded regions represent 95% confidence bands for the correlated response to selection.

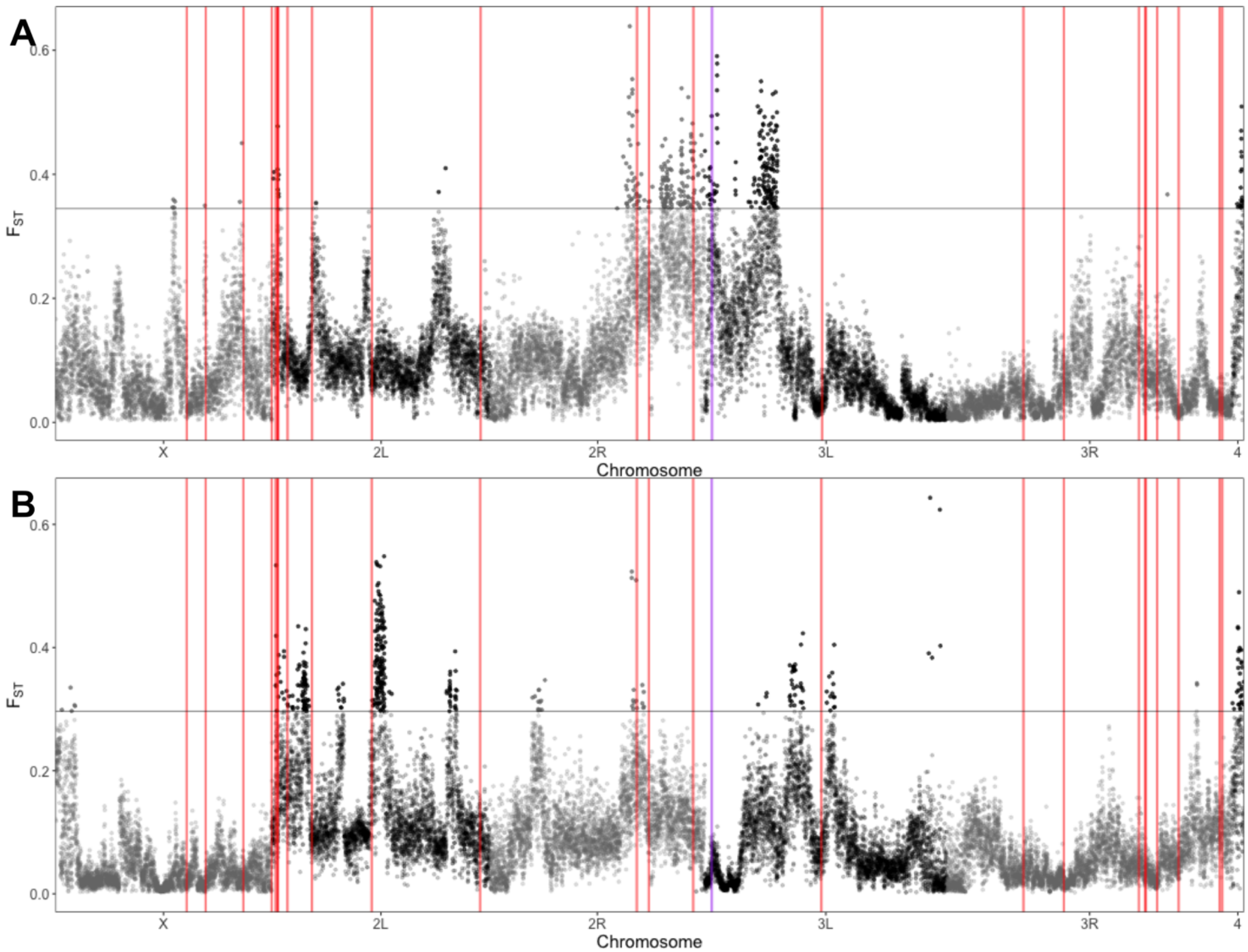


Supplemental Figure 13. Genetic differentiation between artificial selection pools (Figure 3B), with core hippo signaling loci (red) and *emc* (purple) marked. Genomic differentiation between up and down selection lineages ($F_{ST}$) measured in 5000bp windows for the artificial selection along *ds* (A) and *emc* shape change vectors (B). Horizonal grey lines represents 3sd from mean $F_{ST}$.

```
195     GGATTATATGGGGTTTACTAGAAGTGTATATGACCCATCTTAGTTGGCATATTTGTGCTT   108
28      GGATNATATGGGGTTTACTAGAAGTGTATATGACCCATCTTAGTTGGCATATTTGTGCTT   106
96      GGATNATATGGGGTTTACTAGAAGTGTATATGACCCATCTTAGTTGGCATATTTGTGCTT   106
48      GGATNATATGGGGTTTACTAGAAGTGTATATGACCCATCTTAGTTGGCATATTTGTGCTT   106
59      GGATTATATGGGGTTTACTAGAAGTGTATATGACCCATCTTAGTTGGCATATTTGTGCTT   109
801     GGATTATATGGGGTTTACTAGAAGTGTATATGACCCATCTTAGTTGGCATATTTGTGCTT   108
ref     ggattatatggggtgtacaagaagtgtatatgactcatcttag-----------------   163
129     GGATTATATGGGGTGTACTAGAAGTGTATATGACTTATCTTAG-----------------   102
301     GGATTATATGGGGTGTACTAGAAGTGTATATGACTTATCTTAG-----------------   100
69      GGATTATATGGGGTGTACTAGAAGTGTATATGACTTATCTTAG-----------------   102
385     GGATTATATGGGGTGTACTAGAAGTGTATATGACTTATCTTAG-----------------   100
75      GGATTATATGGGGTGTACTAGAAGTGTATATGACTCATCTTAG-----------------   101
83      GGATTATATGGGGTGTACTAGAAGTGTATATGACTTATCTTAG-----------------   100
491     GGATTATATGGGGTGTACTAGAAGTGTATATGACTCATCTTAG-----------------   101
34      GGATTATATGGGGTGTACAAGAAGTGTATATGACTCATCTTAG-----------------   99
774     GGATTATATGGGGTGTACAAGAAGTGTATATGACTCATCTTAG-----------------   102
        **** ********* *** ***************  *******

195     AGGTGCATATTTCTAACAAAAATCCGTAACTATTTTGAGTGGCCAAGAAAATTTCAAATT   168
28      AGGTGCATATTTCTAACAAAAATCCGTAACTATTTTGAGTGGCCAAGAAAATTTCAAATT   166
96      AGGTGCATATTTCTAACAAAAATCCGTAACTATTTTGAGTGGCCAAGAAAATTTCAAATT   166
48      AGGTGCATATTTCTAACAAAAATCCGTAACTATTTTGAGTGGCCAAGAAAATTTCAAATT   166
59      AGGTGCATATTTCTAACAAAAATCCGTAACTATTTTGAGTGGCCAAGAAAATTTCAAATT   169
801     AGGTGCATATTTCTAACAAAAATCCGTAACTATTTTGAGTGGCCAAGAAAATTTCAAATT   168
ref     -ttggcatatttctaacaaaaatccgtaactattttgagtgtccaagaaaatttcaaa-t   221
129     -TTGGCATATTTCTAACAAAAATCCGTAACTATTTTGAGTGGTAGAAA-ATTTCAAAT-T   159
301     -TTGGCATATTTCTAACAAAAATCCGTAACTATTTTGAGTGGTAGAAA-ATTTCAAAT-T   157
69      -TTGGCATATTTCTAACAAAAATCCGTAACTATTTTGAGTGGTAGAAA-ATTTCAAAT-T   159
385     -TTGGCATATTTCTAACAAAAATCCGTAACTATTTTGAGTGGTAGAAA-ATTTCAATT-T   157
75      -TTGGCATATTTCTAACAAAAATCCGTAACTATTTTGAGTGGTAGAAA-ATTTCAAAT-T   158
83      -TTGGCATATTTCTAACAAAAATCCGTAACTATTTTGAGTGGTAGAAA-ATTTCAAAT-T   157
491     -TTGGCATATTTCTAACAAAAATCCGTAACTATTTTGAGTGGTAGAAA-ATTTCAAAT-T   158
34      -TTGGCATATTTCTAACAAAAATCCGTAACTATTTTGAGTGTCCAAGAAAATTTCAAA-T   157
774     -TTGGCATATTTCTAACAAAAATCCGTAACTATTTTGAGTGTCCAAGAAAATTTCAAA-T   160
            *************************************    * * * **  *   *

195     TTTTTTTGCGTGTGCTTGGATGTTGTAATTTTGCATTGTTTGGCTAATCTACGTAAATCT   228
28      TTTTTTTGCGTGTGCTTGGATGTTGTAATTTTGCATTGTTTGGCTAATCTACGTAAATCT   226
96      TTTTTTTGCGTGTGCTTGGATGTTGTAATTTTGCATTGTTTGGCTAATCTACGTAAATCT   226
48      TTTTTTTGCGTGTGCTTGGATGTTGTAATTTTGCATTGTTTGGCTAATCTACGTAAATCT   226
59      TTTTTTTGCGTGTGCTTGGATGTTGTAATTTTGCATTGTTTGGCTAATCTACGTAAATCT   229
801     TTTTTTTGCGTGTGCTTGGATGTTGTAATTTTGCATTGTTTGGCTAATCTACGTAAATCT   228
ref     tttttttgcgtgtgcttggatgttgtaatttcgcattgtttggctaatctacgtaaatct   281
129     TTTTTTTGCGTGTGCTTGGATGTTGTAATTTCGCATTGTTTGGCTAATCTACGTAAATCT   219
301     TTTTTTTGCGTGTGCTTGGATGTTGTAATTTCGCATTGTTTGGCTAATCTACGTAAATCT   217
69      TTTTTTTGCGTGTGCTTGGATGTTGTAATTTCGCATTGTTTGGCTAATCTACGTAAATCT   219
385     TTTTTTTGCGTGTGCTTGGATGTTGTAATTTCGCATTGTTTGGCTAATCTACGTAAATCT   217
75      TTTTTTTGCGTGTGCTTGGATGTTGTAATTTCGCATTGTTTGGCTAATCTACGTAAATCT   218
83      TTTTTTTGCGTGTGCTTGGATGTTGTAATTTCGCATTGTTTGGCTAATCTACGTAAATCT   217
491     TTTTTTTGCGTGTGCTTGGATGTTGTAATTTCGCATTGTTTGGCTAATCTACGTAAATCT   218
34      TTTTTTTGCGTGTGCTTGGATGTTGTAATTTCGCATTGTTTGGCTAATCTACGTAAATCT   217
774     TTTTTTTGCGTGTGCTTGGATGTTGTAATTTCGCATTGTTTGGCTAATCTACGTAAATCT   220
        ******************************* ****************************
```

Supplemental Figure 14. Alignment of sanger sequencing of the region containing the *ds* polymorphism from several DGRP lines (line numbers indicated at line starts) to reference sequence (from *Drosophila melanogaster* genome). DGRP lines are indicated with DGRP 195, 28, 96, 48, 59 and 801 predicted to have the insertion, and, DGRP 129, 301, 69, 385, 75, 83, 491, 34 and 774 without. In addition to the insertion in these lines, several other associated SNPs are found linked to the insertion.

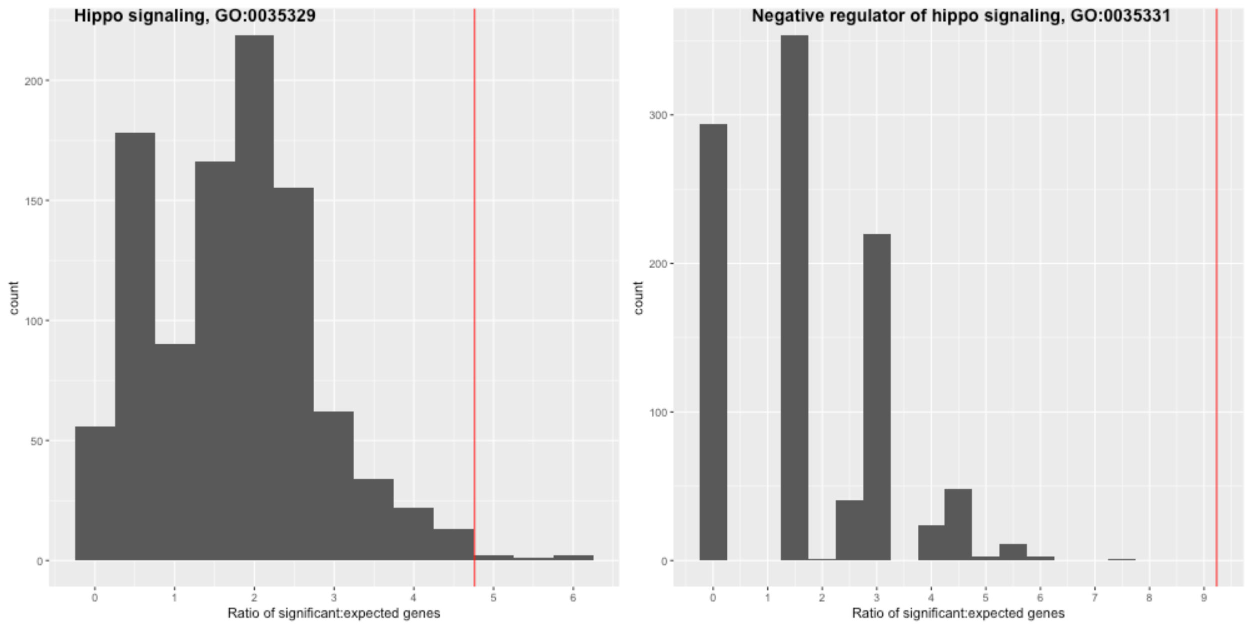


Supplemental Figure 15: Permutation test for over representation of hippo signaling terms in outlier regions. The red line represents the observed ratio of significant to expected genes in outlier windows ($F_{ST}$ greater than 3 standard deviations from the mean). The permutation test selected random windows from the genome, equal in number to those identified as outliers. For each random draw of loci, the ratio of significant to expected genes in the term of interest was calculated for 1000 permutations.

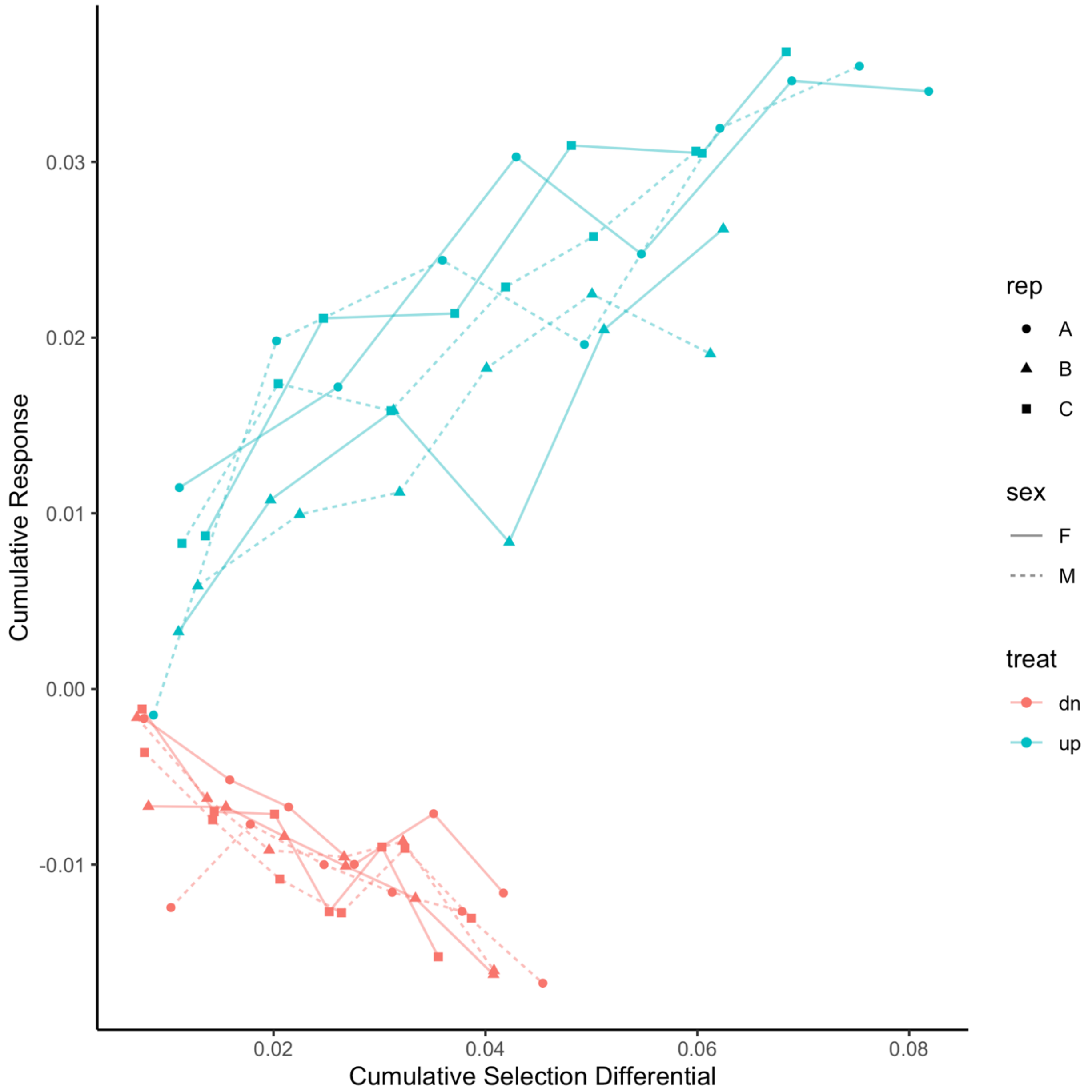


Supplemental Figure 16. Response to selection based on projections onto *emc* shape change vector. Regression of cumulative selection differential onto cumulative response was used to estimate realized heritability in both “up” and “down” selection regimes independently.

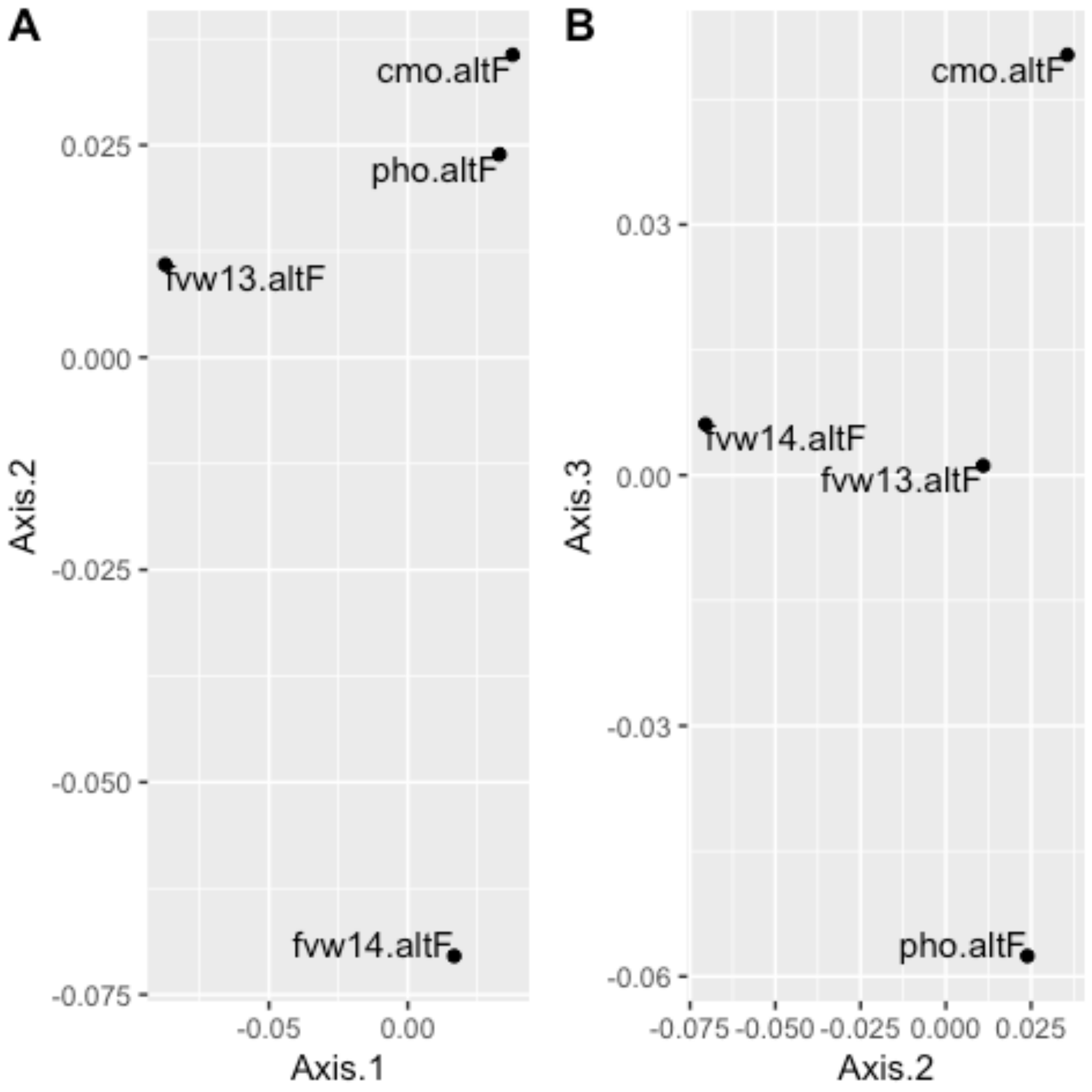


Supplemental Figure 17. Minimal genetic structure among wild populations used in this study. Principal coordinate plot calculated from Bray's distances estimated from allele frequency data (altF), between the four wild cohorts used in this analysis. Axis 1-3 explains 45%, 30% and 25% of the variance in genetic distances respectively.

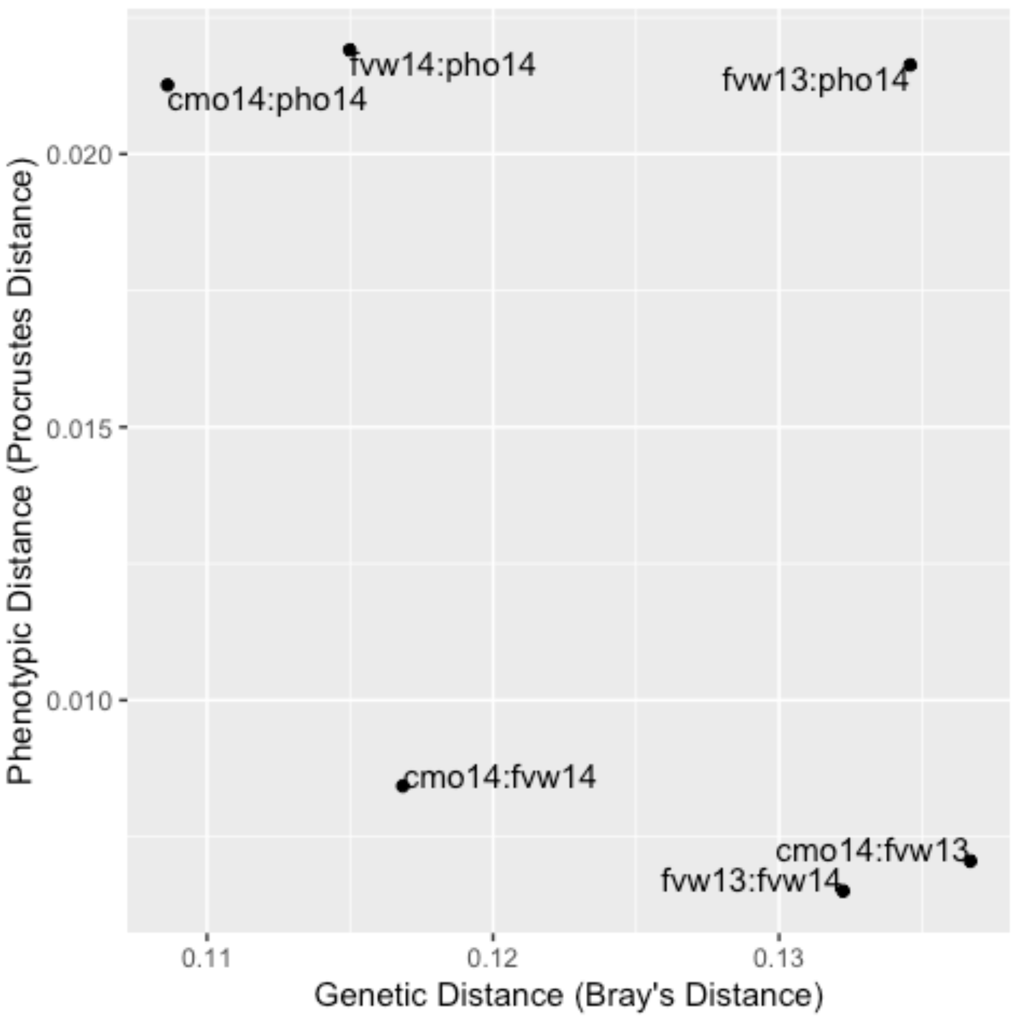


Supplemental Figure 18. Genetic distance is not correlated with phenotypic distance among wild cohorts used in this study. Phenotypic distances are the Euclidian distances from model estimated shape vectors for each population from a multivariate regression of shape onto for population and wing centroid size.

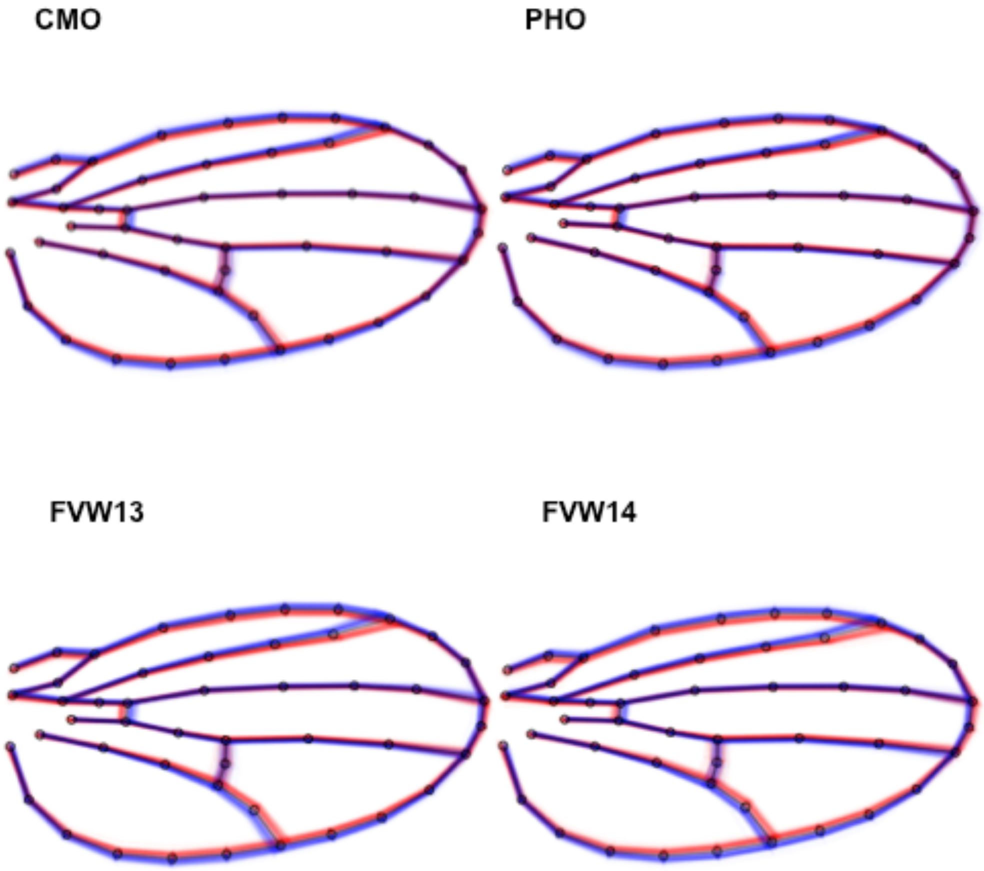


Supplemental Figure 19. Shape variation within *ds* selected pools for bulk segregant analysis by population. Wings within selected pools (one red and one blue, representing the “up” and “down” pools) are plotted to show the phenotypic extremes for the shape scores used for selecting individuals within and between pools. Black line indicates mean shape between pools.

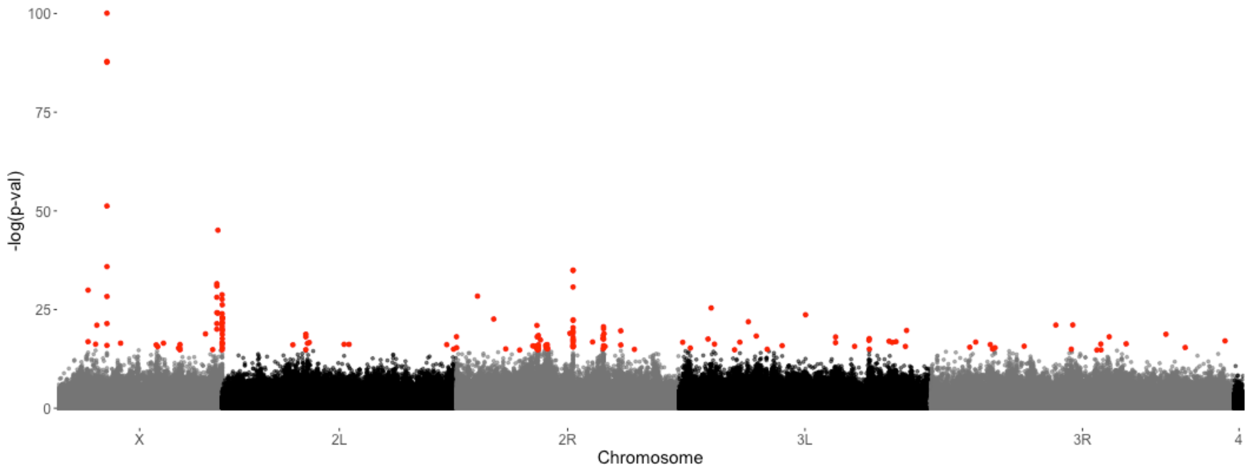


Supplemental Figure 20. Removing PHO from the BSA analysis increases the number of differentiated sites between *ds* shape change pools. Genome-wide scan for differentiated loci between pools selected based on *ds* shape change vector using the CMH test using ACER. Variants in *ds* are still not implicated in this analysis. Points in red indicate sites with significant differentiation based with a FDR of 0.05.

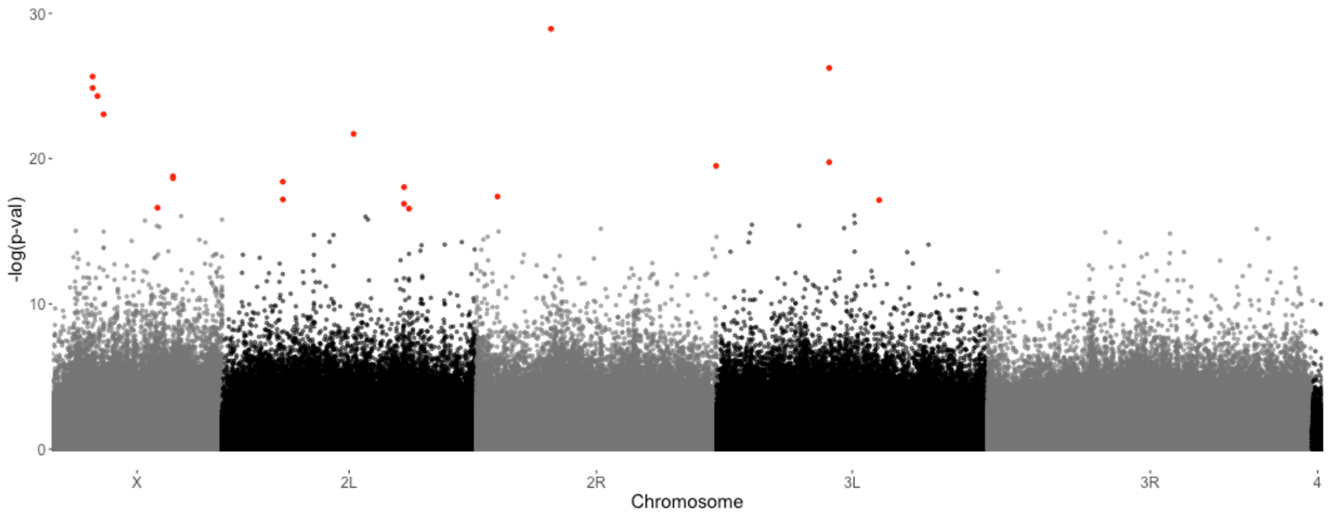


Supplemental Figure 21. Down sampling genome coverage does not impact results for CMH tests substantially. When all pools are sampled to a coverage depth of 75 reads, while preserving allele frequency, we do not find an increase in number of differentiated sites. Although the sites identified as significantly differentiated do change somewhat, in part due to sampling procedure as any sites without a minimum depth of 75 reads in each of the 4 populations was dropped from this analysis. Variants in *ds* are still not implicated in this analysis.

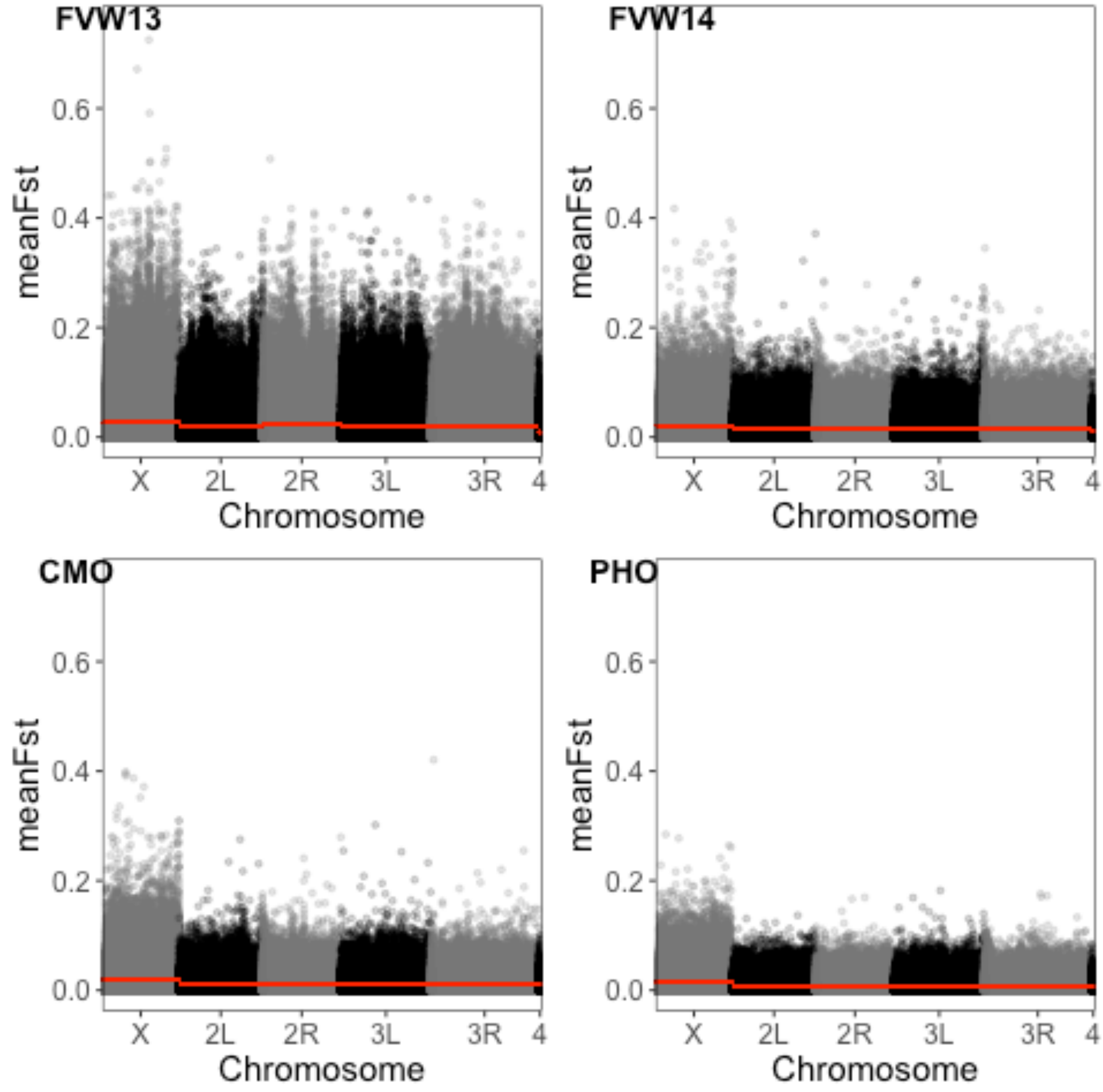


Supplemental Figure 22. $F_{ST}$ between pools of individuals selected along the *ds* shape change axis within each population. Calculated in 100 bp windows using PoPoolation2 program. Elevated $F_{ST}$ on the X chromosome is due to sampling of fewer X chromosomes, relative to autosomes as most pools, with the exception of FVW14, consist of only males. Red line indicates the mean $F_{ST}$ for the chromosome, which as expected is very low.

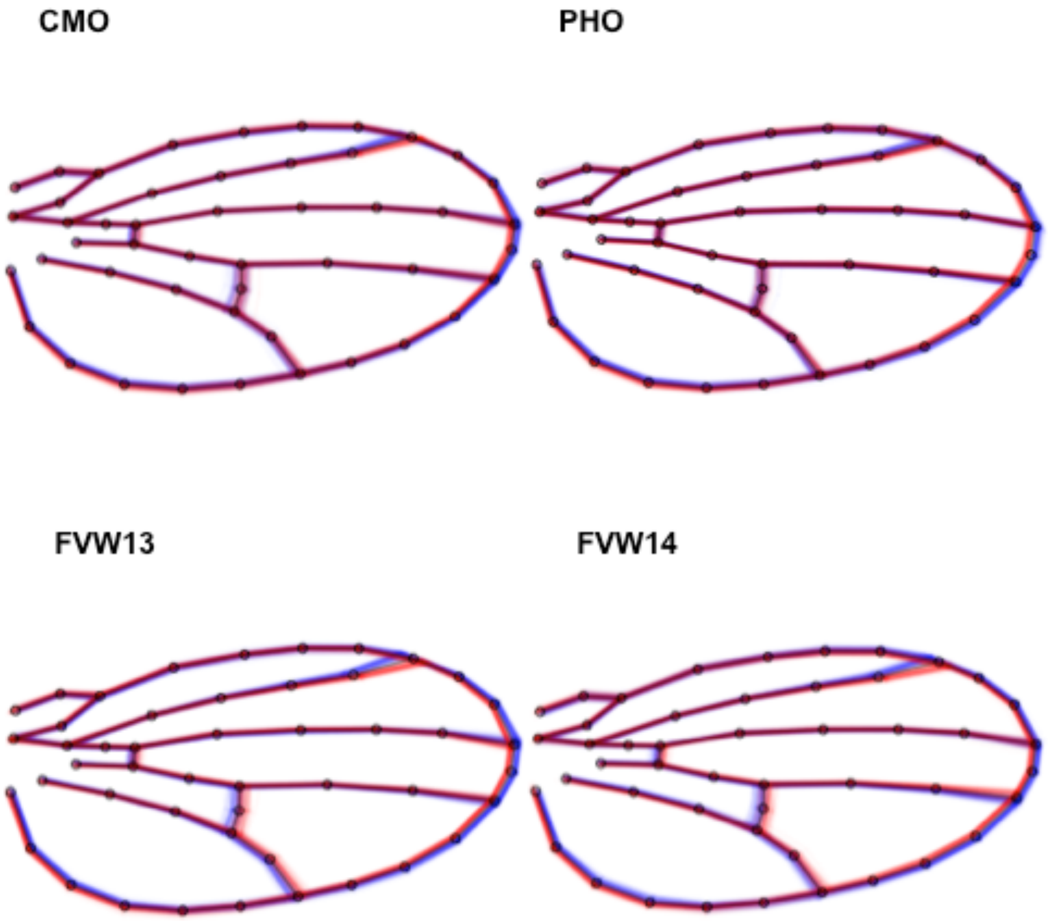


Supplemental Figure 23. Shape variation within *neur* selected pools for bulk segregant analysis by population. Wings within selected pools (one red and one blue, representing “up” and “down” extreme pools respectively ) are plotted.

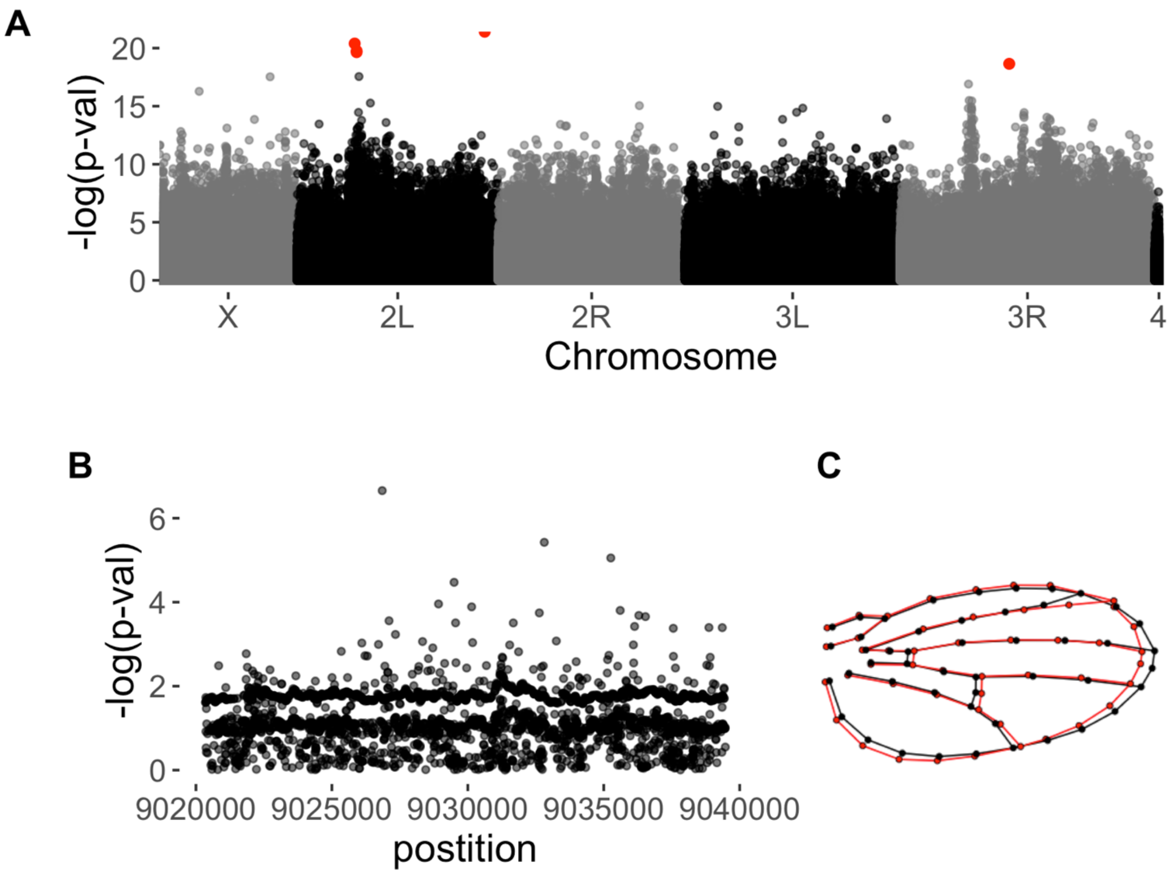


Supplemental Figure 24. Genome-wide scan for differentiated loci between pools selected based on *neur* shape change vector using the CMH test implemented with ACER. (A) Whole genome scan for differentiation. Points in red indicate sites with significant differentiation based with a FDR of 0.05. (B) No significantly differentiated sites within *neur.* (C) shape change between selected pools based on *neur* shape change vector, effect size is multiplied by 2 for visualization.

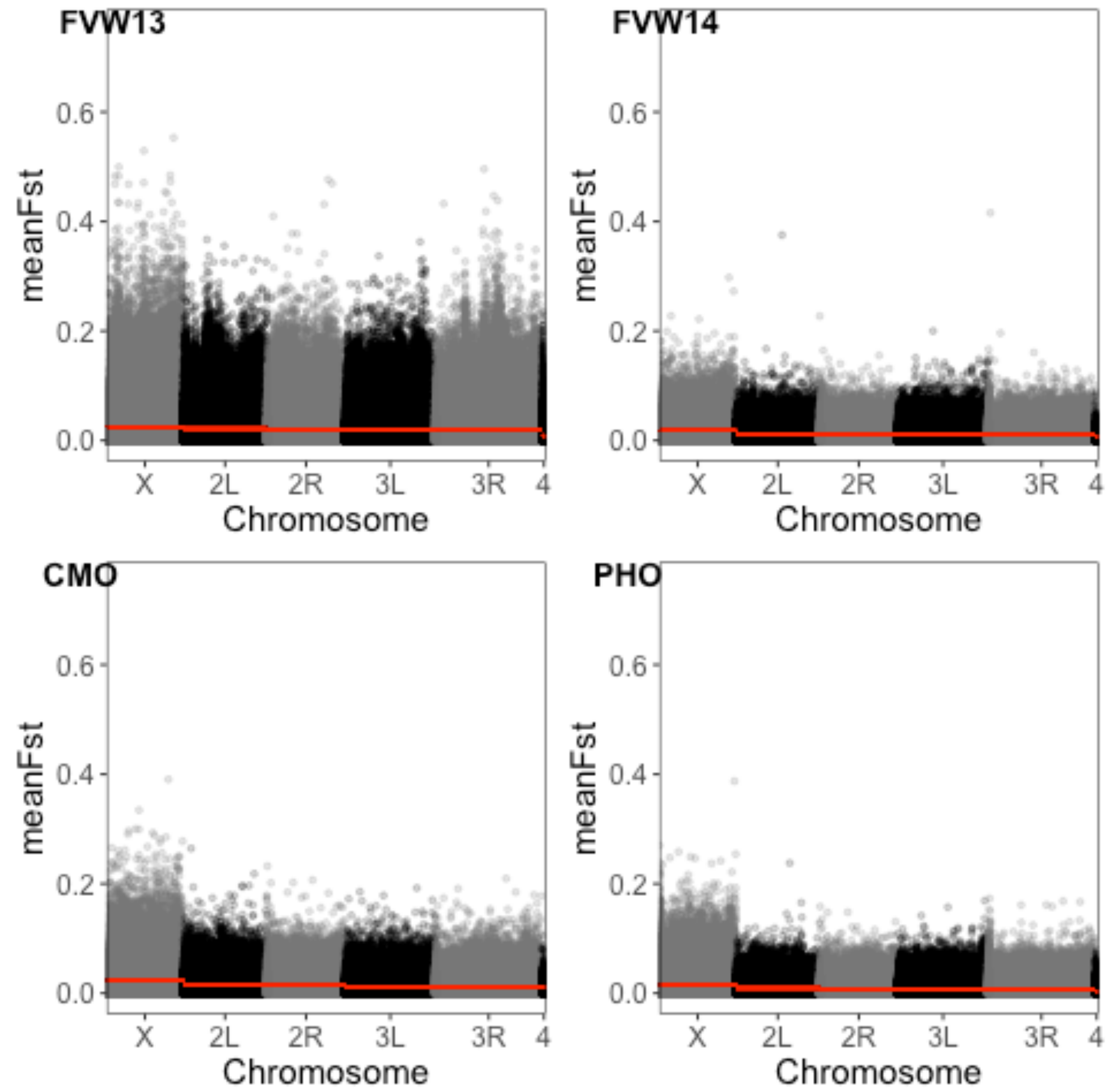


Supplemental Figure 25. $F_{ST}$ between pools of individuals selected along the *neur* shape change axis within each population. Calculated in 100 bp windows using PoPoolation2 program. Elevated $F_{ST}$ on the X chromosome is due to sampling of fewer X chromosomes, relative to autosomes as most pools, with the exception of FVW14, consist of only males. Red line indicates the mean $F_{ST}$ for the chromosome.
